# CIA5 INTERACTS WITH THE ZINC CHAPERONE ZNG3 TO BALANCE CARBON AND ZINC METABOLISM

**DOI:** 10.1101/2025.08.16.670667

**Authors:** George Kusi-Appiah, Stefan Schmollinger, Andrew Mamo, Sarah C. Stainbrook, Thomas V. O’Halloran, Daniela Strenkert

## Abstract

Carbon and zinc (Zn) metabolism are intrinsically connected in phototrophs, as crucial components involved in CO_2_ assimilation, like carbonic anhydrases, are highly abundant Zn proteins. Utilizing these and other proteins, the eukaryotic green algae *Chlamydomonas reinhardtii* can maintain phototrophic growth in low CO_2_ environments by inducing a carbon concentrating mechanism (CCM). In this work we show that Chlamydomonas dynamically increases its Zn content to accommodate the higher intracellular Zn demand in low CO_2_ environments. This increase requires the presence of Cia5, a major regulator of the CCM in Chlamydomonas. How Cia5 regulates expression of thousands of low CO_2_-inducible genes remains enigmatic, its transcript and protein abundance is unchanged in different CO_2_ environments, even in the presence of an additional reduced carbon source, acetate. We show here that the Cia5 protein is not present in Zn-limitation, despite *CIA5* transcription being unchanged. We used a CRISPR knock-in approach to express Cia5-HA from its endogenous locus and used two independent Cia5-HA expressing strains for affinity purification and identified a protein belonging to a conserved family of metal binding GTPases, ZNG3, as a constitutive interaction partner. Like Cia5, ZNG3 is constitutively expressed, co-expressed with Cia5 along the diurnal cycle and is Cia5-dependently induced in low CO_2_ environments. Surprisingly, *zng3* mutants do not phenocopy *cia5* mutants and grow well in low CO_2_ conditions. Instead, *zng3* mutants are unable to grow like wildtype if excess carbon is available in the form of high CO_2_ or acetate. Transcriptomics of wildtype and *zng3* mutants grown with different carbon sources revealed that transcriptional induction of the majority of genes involved in the CCM is maintained in low CO_2_ grown *zng3* mutants, while the degree of induction in a subset of *LCI* genes is reduced (*HLA3*, *CAH4* and *CAH5*). Genes encoding proteins involved in plastid quality control were induced in *zng3* mutants grown on acetate and high CO_2_, as well as other, related metallochaperones. We hypothesize that Zn trafficking towards the plastid is mis regulated in *zng3* mutants resulting in protein mis-metalation and unfolding. Taken together, we propose that ZNG3 and Cia5 coordinate Zn and CO_2_ metabolism, affecting intracellular Zn trafficking and modulate the CO_2_ response.

## INTRODUCTION

Algae are crucial primary producers, accounting for ∼50% of all photosynthetic carbon fixation on earth [1, 2], and are equally important as model organisms for fundamental research in photosynthesis and nutrient metabolism [3]. Algae thrive in diverse ecological niches not used for agriculture, even in environments that are characterized by extreme conditions [4], demonstrating substantial metabolic flexibility and adaptability [5].

Many eukaryotic and prokaryotic algae, including the tractable model *Chlamydomonas reinhardtii* (Chlamydomonas hereafter), utilize carbon concentrating mechanisms (CCM) to grow efficiently in their naturally low CO_2_ habitats [6–9]. The CCM facilitates the uptake of both HCO_3_^-^ and CO_2_ against a gradient, achieving ∼10-40 fold higher inorganic carbon (C_i_) concentrations intracellularly compared to what is found in their direct, natural environment [10]. Transporters for CO_2_/HCO_3_^-^ mediate cellular or organellar import [9] and carbonic anhydrases (CAHs) facilitate interconversion of CO_2_ to HCO_3_^-^, either to aid transport or provide the CO_2_ substrate for fixation [11–15]. Ribulose-1,5-bisphosphate (RuBP) carboxylase-oxygenase (RuBisCO), at the center of the Calvin-Benson cycle in the chloroplast, is ultimately where CO_2_ is assimilated onto RuBP, producing two molecules of 3-phosphoglycerate (3PGA) [16]. RuBisCO also incorporates O_2_ onto RuBP, the rate of which depends on temperature, CO_2_:O_2_ ratio and inherent enzymatic properties, ∼3 CO_2_ to 1 O_2_ are assimilated in ambient conditions [17]. RuBP oxygenation produces one molecule of 2-phosphoglycolate (2PG) instead of one 3PGA, which then needs to be costly (CO_2_ loss, additional ATP and NAD(P)H investment) recycled via photorespiration [18]. RuBisCO selectivity (CO_2_:O_2_) can be improved by increasing the CO_2_ concentration and minimizing O_2_ exposure around the enzyme. For this purpose eukaryotic algae such as Chlamydomonas use pyrenoids [11], which are self-assembling, membrane-less structures that utilize starch sheets to create a diffusion barrier around RuBisCO [19]. Similar strategies are found in diatoms that use protein-walled pyrenoids [20], while cyanobacteria fix CO_2_ in bacterial protein microcompartments called carboxisomes [21], and land plants developed C_4_ [22] and CAM metabolism [23] to decrease CO_2_ loss in photosynthetic carbon fixation [24].

Chlamydomonas’ CCM is not expressed constitutively but dynamically induced when cells are exposed to very low, or air levels of CO_2_ (0.04%) during growth [10]. A forward genetic screen identified a crucial regulator of the CCM, the constitutively expressed Cia5/CCM1 (Cia5 hereafter) [25–27]. *cia5* mutants fail to transcriptionally induce C_i_ transporters, CAHs and many structural components of the CCM, and accordingly have a growth defect with low CO_2_ supply [25, 26, 28, 29]. Global transcriptomic studies identified many more low-CO_2_-inducible (*LCI*) genes that fail to respond either partially or fully in *cia5* mutants, indicating a wider restructuring of Chlamydomonas metabolism in response to low CO_2_ [28]. Complementation studies identified two crucial regions for Cia5 function, ∼ 110 amino acids at the N-terminus and ∼130 amino acids towards the C-terminus were sufficient to restore growth in very low CO_2_ conditions and partially restored molecular phenotypes [30]. A second regulator, LCR1, downstream of Cia5, was identified to be involved in regulating a subset of genes involved in low CO_2_ acclimation [31].

It is still unclear mechanistically how Cia5 is activated to induce the Chlamydomonas CCM, but previous work highlighted the dependence of successful acclimation to low CO_2_ environments on Zinc (Zn) availability [32, 33]. Zn is a common cofactor in eukaryotes, ∼4-10% of all proteins are estimated to utilize Zn as an essential cofactor. While the mechanisms of over four hundred eukaryotic enzymes depend upon catalytic steps mediated by Zn in the active site, the majority of Zn-dependent proteins, mostly utilize Zn-binding to, mostly to stabilize protein structure for example in Zn finger domains that facilitate DNA, RNA or protein interactions [34–37]. One example is the CCM regulator Cia5, which contains two Zn binding sites within the first, 110 amino acids that are essential for function [34, 35]. The first *cia5* mutant identified indeed had a single amino acid change affecting the first Zn binding sites (H54Y) [25–27]. BSD2 is a Zn finger protein requiring two Zn atoms, localized to the chloroplast involved in CO_2_ assimilation, [38].

BSD2 is a small, RuBisCO-specific chaperone involved in the assembly of its L_8_S_8_ hexadecameric form, and required to allow CO_2_ fixation [38–40]. CAHs are abundant and integral enzymes to CCMs, present in all cellular compartments and they do rely on Zn in their catalytic center for the reversible hydration of CO_2_ to HCO_3_^-^ [41, 42]. In marine habitats, where Zn can be scarce [43], cobalt (Co) can replace Zn in catalytic centers of some CAHs, or backup enzymes that rely on cadmium (Cd) can be used by some organisms instead [44, 45]. That such Zn-sparing mechanisms evolved underscores the crucial contribution of CAHs and Zn to the phototrophic lifestyle [45, 46]. Transporters involved in Zn import and distribution as well as shuttles have been identified in Chlamydomonas using transcriptome and proteome studies in low Zn environments, but it is still yet unclear how Zn homeostasis is integrated into larger cellular regulatory networks [33, 47, 48].

In this work we identified ZNG3, a protein belonging to the COG0523/CobW family of Zn chaperones [49], directly and condition-independently interacting with Cia5 *in vivo*. Proteins belonging to the family of Zn regulated GTPases (ZNG) have been identified as candidate Zn chaperones that are capable to transfer Zn to Zn proteins based on domain structures, Zn binding capabilities and GTPase activity [50]. While nuclear localized Cia5 contains a Cys_2_His_2_ (C_2_H_2_) zinc-finger domain, it also contains a second, unusual potential Zn binding site, both of which may be targets for metalation by ZNG3. While it has been shown that ZNG1 proteins interact with their clients via the Zn finger domain, Zn transfer seems to be mediated towards another Zn binding site within the client protein [51]. Because of this, it is likely that ZNG3 delivers Zn to the second Zn binding site within Cia5. We propose that ZNG3 and Cia5 form a bipartite, regulatory module that enables cells to promptly and dynamically adjust their Zn and carbon status in response to changes in CO_2_ availability in their environment. ZNG3 seems to be crucial for blocking Zn trafficking towards the plastid when no CCM is established and thus Zn demand is lower, as *zng3* mutants maintain relatively high Zn levels in this situation. We show that excess Zn is concomitant with the induction of the plastid unfolded protein response in *zng3* mutants suggesting protein mis metalation. We suggest that Cia5 activity *per se* is not dependent on ZNG3 as *zng3* mutants are asymptomatic when grown in low CO_2_ conditions.

## RESULTS

### CO_2_ assimilation and Zn metabolism are connected

Chlamydomonas’ ability to induce the CCM in low CO_2_ environments and therefore maintain high phototrophic growth rates is significantly impaired when Zn becomes co-limited [32]. This suggested that the function of one or more Zn-dependent components of the CCM is essential for low CO_2_ acclimation, with abundant Zn-containing proteins like carbonic anhydrases (CAHs) being likely candidates. To assess whether the induction of the CCM, which requires a large increase in Zn-containing protein abundance is impactful enough to require substantial changes in intracellular Zn demand, we measured changes in the Zn content of cells grown with and without additional CO_2_ supplementation. Since expression of many high abundant Zn containing proteins involved in the low CO_2_ response is dependent on the regulator Cia5, we also generated *cia5* mutants to assess if observed changes are also Cia5 dependent. We used a previously established CRISPR/Cpf1 protocol [52, 53] in which we provide a single stranded DNA repair template (ssODNs) and independent guide RNAs to introduce two *in-frame* stop codons into the first exon of the *CIA5* gene, and subsequently identified two independent strains that carried the respective insertions (Figure 1A). Loss of Cia5 function was validated by probing the expression of a known target of Cia5 which is required for successful acclimation to low CO2, the mitochondrial β-CA CA4 (*CAH4*) [28, 54, 55]. Transcripts for *CAH4* were highly induced in wildtype in low CO_2_ (0.04%, LC) phototrophically grown cultures, but not in the two *cia5* mutants, (Figure 1B). As expected, neither wildtype nor *cia5* mutants showed *CAH4* expression with high CO_2_ (5%, HC) supplementation, consistent with previous reports [28]. We then analyzed the Zn content in wildtype and *cia5* mutants, both in photoheterotrophically (acetate as additional carbon source, Ac) and phototrophically (HC and LC) grown cultures using Inductively Coupled Plasma Tandem Mass Spectrometry (ICP MS/MS). Phototrophically grown, CO_2_ limited (LC) wildtype cells increased their Zn quota (the sum of all Zn atoms in a cell) by ∼70% (p<0.5) as compared to cultures supplemented with CO_2_ (HC), or by ∼ 50% (p<0.5) compared to cultures grown photoheterotrophically (Figure 1C). Cellular quotas for other metals including Fe and Mn do not change to a significant degree under these conditions (Supplemental Figure 1). The increase in Zn content was not observed in low CO_2_ grown *cia5* mutants, consistent with a lack of acclimation to low CO_2_ environments in these mutants and loss of CAH expression (Figure 1C).

The increase of the Zn quota in LC grown wildtype cultures reflects the requirement of relatively high abundant, Zn-containing proteins expressed conditionally in low CO_2_. Since this increase upregulated in LC are a part of this increase. The increase in Zn in LC further highlights the impact of carbon availability on Zn metabolism. We expect that a substantial increase in the Zn quota of Chlamydomonas cells during CCM establishment necessitates elaborate adjustments to the cell’s Zn metabolism, including induction of Zn importers and Zn handling proteins.

**Figure 1.**
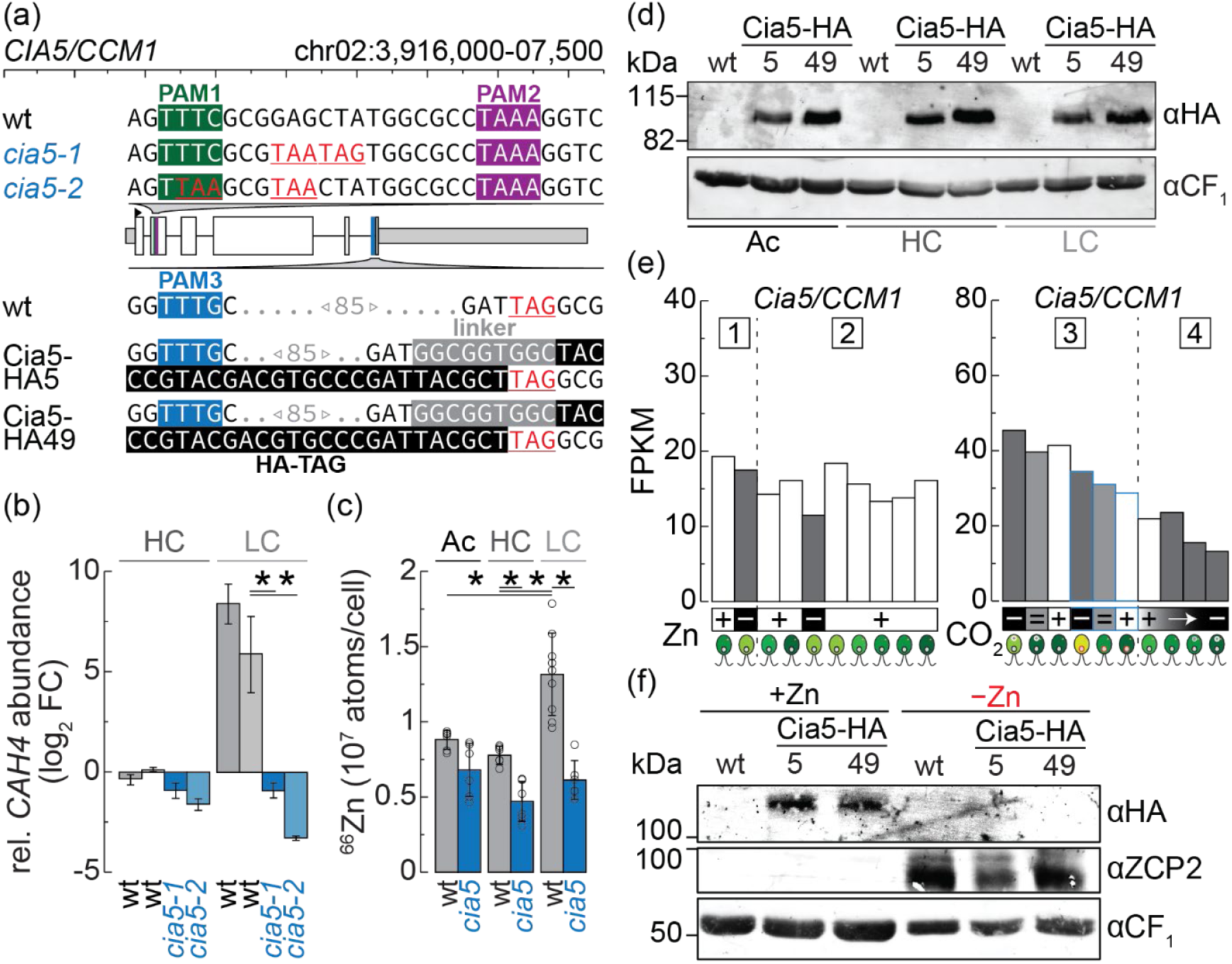
CO_2_ and Cia5-dependent adjustment of the Zn quota in Chlamydomonas, Cia5 is constitutively expressed independent of the carbon source but not present in Zn deficient cells. (a) Illustration of the *Cia5* locus with the utilized PAM sites to generate CRISPR mutants (green and magenta) or HA knock-ins (blue). Introduced *in-frame* stop codons in *cia5-1* and *cia5-2* are highlighted red, introduced sequences coding for a short linker (grey) and HA epitope (black) are shown for Cia5-HA5 and Cia5-HA49 before the native stop (red) at the C-terminus. (b) Transcript abundances of *CAH4* were determined using qRT-PCR in wildtype and *cia5* mutants grown phototrophically with supplementation of either 5% CO_2_ (HC) or 0.04% CO_2_ (LC). Averages and standard deviation of 3 independent cultures are shown as log_2_ fold changes compared to wildtype in HC, asterisks indicate significance (two-sided T-test, p < 0.05). (c) Wildtype (wt) and *cia5* mutants were grown in HC, LC and photoheterotrophically (Ac, with acetate as a carbon source). Zn content was determined by ICP-MS/MS and normalized to cell numbers. Shown are averages and standard deviation of 3-9 independent experiments, as well as individual data points, asterisks indicate significance (two-sided T-test, p < 0.05). (d) Total protein was extracted from wildtype (wt) and both Cia5-HA lines grown in Ac, HC or LC conditions. Denatured proteins were separated with SDS-PAGE, transferred to nitrocellulose and immunodecorated with antibodies against the HA epitope, antibodies specific to CF_1_ served as loading control. (e) Survey of transcript abundances of *CIA5* in published RNAseq datasets with varying Zn and CO_2_ supply. Malasarn *et al.* (1) analyzed transcripts in Zn-replete (+) and Zn-deficient (−) cultures, Hong Hermesdorf *et al*. (2) analyzed early exponential and early stationary Zn-replete cultures (+), as well as Zn-deficient (−) cultures and Zn resupply. Fang *et al.* analyzed transcript abundances in cultures acclimated to high (+, 5%), air-level (=, 0.04%) and low CO_2_ (−, 0.01%) in wildtype (black outline) and *cia5* mutants (blue outline), while Brueggemann *et al*. (3) analyzed the transition from high (+, 5%) to very low CO_2_ supply (−, 0.01%) (f) Total protein was extracted from wildtype and both Cia5-HA lines, grown photoheterotrophic with (+) or without (−) Zn supplementation. Denatured proteins were separated with SDS-PAGE, transferred to nitrocellulose membranes and decorated with antibodies specific to the HA epitope. Antibodies specific to CF_1_ served as loading control, ZCP2 served as control for successful establishment of Zn deficiency.

### Constitutive Cia5 expression is Zn dependent

To probe the expression of Cia5 protein and identify potential interacting proteins, we used our CRISPR/Cpf1 gene editing approach to introduce an HA tag to the C-terminus of the native *CIA5* locus (including a small, flexible glycine linker, Figure 1A). We identified two independent strains that express the Cia5-HA fusion protein from their endogenous locus and that harbored no unintended mutations or frameshifts, named Cia5-HA5 and Cia5-HA49 (Figure 1AD). The CRISPR/CPf1 knock-in approach has the advantage that *CIA5*’s endogenous gene regulation is maintained and that only a single, HA-tagged copy of Cia5 is present in the cell. We selected a commercial anti-HA antibody that is sensitive and has no unspecific cross-reactivity when used on total cell lysate of Chlamydomonas. This allowed us to monitor Cia5 protein abundance under different conditions, including when cells were grown phototrophically (HC/LC), or photoheterotrophically (Ac). We validated earlier reports [25] that Cia5 is constitutively expressed in phototrophically grown cultures independent of CO_2_ supplementation (Figure 1D), and additionally found that Cia5 is equally unchanged when cells are grown photoheterotrophically (Figure 1D). Since Cia5 is constitutively expressed and localized to the nucleus independent of CO_2_ availability [25, 26, 28, 29], it was proposed previously that posttranslational modifications may mediate its modular, regulatory role in establishing the CCM. In this study, we did not observe a noticeable shift in protein size or multiple bands in any condition. We next surveyed *CIA5* transcript abundance in previously published transcriptome datasets probing alterations in Zn or CO_2_ metabolism [28, 47, 56, 57]. Malasarn *et al.* analyzed transcript abundance in photoheterotrophic Zn-replete and Zn deficient cultures [56], while Hong-Hermesdorf *et al.* compared Zn replete cultures from early logarithmic and beginning stationary stages with Zn deficient cultures and further monitored recovery of transcript abundances after the addback of Zn to those Zn-deficient cultures in a time-dependent manner (0, 1.5, 3, 4.5, 12, 24h, Figure 1E, left panel). Fang *et al.* analyzed RNA abundance in wildtype and the *cia5* mutant (original mutant H54Y, with a point mutation in the histidine residues of the first C_2_H_2_ Zn binding site) cultures acclimated high CO_2_ (5%), air-levels of CO_2_ (0.04%) and low CO_2_ (0.01 %) [28], while Brueggemann *et al.* followed the transition of wildtype cultures from high (5 %) to low (0.01 %) CO_2_ conditions in a time course (0, 0.5, 1, 3h, Figure 1E, right panel) [57]. *CIA5* transcript abundance did not show a significant increase or decrease in response to changes in Zn or CO_2_ nutrition in any of these experiments, but we did note a minor, transient increase of *CIA5* transcripts upon Zn resupply to Zn-deficient cultures (Figure 1E). Because Cia5 is a Zn containing protein, we probed Cia5 protein abundance in Zn replete and Zn deficient conditions using the strains that express the Cia5-HA fusion proteins (Figure 1F). We used expression of ZCP2 as a marker protein for Zn deficiency [33, 56] to validate successful Zn exclusion from the growth media and induction of the Zn deficiency response. ZCP2 is a predicted but uncharacterized Zn chaperone that is expressed solely in the absence of Zn from the growth media [33], an expression profile that we recapitulated in our experiment (Figure 1F). As expected, Cia5-HA accumulated in Zn replete conditions in both Cia5-HA expressing strains (5 and 49), but no Cia5-HA was detected in cells grown in Zn limitation (Figure 1F). While Cia5 is a Zn-finger protein with an additional Zn binding site, this was still surprising, since *CIA5* transcript abundance is unchanged when cells are grown in Zn deficiency. In addition, Cia5 is not a highly abundant protein and would thus not require a large fraction of the cellular Zn pool, which is why it was assumed that regulatory proteins like Cia5 may be unaffected in Zn limited cells [56].

Given that *CIA5* is constitutively transcribed independent of CO_2_ or Zn supply, it is striking that the Cia5 protein accumulates in Zn-replete but not under Zn-deficient growth conditions. This data highlights the dependence of successful CCM induction on Zn availability, utilizing protein stability or translational control of Cia5 to prevent induction of a metabolic pathway for which the required resources are not available.

### Cia5 constitutively interacts with a Zn-binding GTPase

Given that there are no obvious changes in Cia5 protein size or abundance when cells are grown in different, relevant growth conditions (HC/LC/Ac) we assumed that Cia5 dependent establishment of the CCM may be realized by modular protein-protein interactions instead. A likely scenario would be that distinct, condition specific interaction partners can dynamically shape the cell’s response to changes in Zn and CO_2_ availability. To capture potential binding partners of Cia5, we performed immunoprecipitations (IPs) on cell lysates from the two independent Cia5-HA expressing strains grown under LC, HC or Ac conditions using commercially available antibodies targeting the HA epitope. IPs on wildtype strains that do not express an HA-fusion protein served as a negative control and allowed us to identify unspecific contaminants in our experiment. Proteins enriched in IPs were identified by LC-MS/MS using the Chlamydomonas genome v6 as a reference [58]. We only considered proteins as being high confidence interaction partners of Cia5 if they were identified with at least two peptides in IPs from both Cia5-HA5 and Cia5-HA49, but not in wildtype samples in any condition. A total of 40 proteins met these criteria in HC, LC and Ac, respectively (Supplemental Data 1, Figure 2A).

Two proteins were found in all three conditions, while 19 were exclusive to phototrophic conditions and 21 were found exclusively in photoheterotrophic conditions. Some immunoprecipitated proteins that met our criteria may still represent unspecific contaminants rather than Cia5-interacting proteins, which is why we also used the precursor ion intensity as an indicator for the abundance of these proteins in the samples. Two of the obtained proteins were immunoprecipitated in all growth conditions far more abundantly than the others (at least 10 times more enrichment in IPs as compared to all other proteins, Figure 2B) and were the same two that were found in all three conditions. The first one was the bait protein Cia5-HA itself, validating that the IPs were successful in enriching Cia5 from every growth condition. The second protein was a protein of unknown function, containing a nuclear localization signal (Supplemental Figure 2) and a CobW domain belonging to a multi protein family of candidate Zn handling GTPases (hereafter depicted as ZNG3). Based on the large ZNG3 abundance in our Cia5-HA IPs and the potential relevance of this protein of unknown function in Zn metabolism, we decided to functionally characterize ZNG3.

### ZNG3 belongs to a conserved class of Zn binding GTPases

Phylogenetic analyses revealed that ZNG3 belongs to the G3E family of P-loop GTPases, that contain a CobW, also referred to as COG0523, domain [49]. A subgroup among this family, nucleotide-dependent metallochaperones (NMCs), can be distinguished by the presence of a metal binding motif (GCxCC) in between the Switch I and Walker B motifs of the CobW domain. A multiple sequence alignment of ZNG3 with *Arabidopsis thaliana* ZNG1 and 2 *Saccharomyces cerevisiae* ZNG1 [50, 51, 59], and Chlamydomonas proteins containing the CobW domain [60] revealed the presence of the metal binding motif in ZNG3 (Supplemental Figure 3), along with a good conservation of all other canonical motifs. This suggests that ZNG3 belongs to the subgroup of CobW proteins that contain the conserved N-terminal GTPase domain that is implicated in metal transport to client proteins via GTP hydrolysis [49, 61]. Some of the CobW proteins, conserved between prokaryotes and eukaryotes, are predicted to be involved in regulating Zn homeostasis due to their induction in Zn limitation [49, 62, 63]. We therefore surveyed *ZNG3*, and other CobW-domain containing proteins in Chlamydomonas, Supplemental Figure 3) [47, 56]. In total, the Chlamydomonas genome encodes 11 proteins with a CobW domain, 3 more containing only the CobW C portion (Supplemental Figure 3). *ZNG3* transcription is independent of the Zn nutritional status of the cell as *ZNG3* transcripts accumulated similarly in Zn-replete and in Zn-deficient conditions (Figure 2C). We did, however, note a minor, transient increase in *ZNG3* abundance upon Zn addback to Zn-deficient cultures, similar to what we have observed for *CIA5* mRNA abundance in the same experiment (Figures 1E and 2C). We probed abundance of the corresponding ZNG3 polypeptide by reanalyzing a previously published proteome from cells grown either in Zn replete or in Zn limited conditions [48] (Figure 2D) and by performing immunodetections using ZNG3 specific peptide antibodies on samples from Zn replete and deficient cells (Figure 2E). ZNG3 protein abundance was indeed more than 14-fold reduced in the proteome of Zn deficient cells as compared to Zn replete grown cells (Figure 2D). Our immunodetections confirmed the reduction of ZNG3 protein when Zn was limited, as ZNG3 protein was below detection limit in a Zn deficient Chlamydomonas culture extract (Figure 2DE). We used our ZNG3 antibodies to perform additional IP experiments from Chlamydomonas cultures using ZNG3 as bait and *zng3-1* as a negative control and indeed identified Cia5 interacting with ZNG3 (Supplemental Data 2). We also analyzed datasets with regards to CO_2_ nutrition for the transcript accumulation of *ZNG3* (Figure 2C). *ZNG3* transcripts increasingly accumulated in low CO_2_ grown wildtype Chlamydomonas cultures, but not in *cia5* mutants and in cells that transition from high to low CO_2_ conditions (Figure 2C, right panel). This indicates that *ZNG3* is a transcriptional target of Cia5, that has some base expression but is further induced upon activation of the CCM. Among Chlamydomonas CobW proteins, ZNG3 expression was most similar to Cre16.g692901, which was the only other CobW protein whose transcripts were not affected by Zn but were Cia5-dependently inducted in response to limiting CO_2_ (Supplemental Figure 3). *ZCP1/2* were solely induced in low Zn, *ZNG2* and *Cre09.g801069* responded both to low CO_2_ (Cia5-dependently) and low Zn, Cre16.g658000 was the only CobW protein whose transcripts were repressed by Zn, while 4 other genes, *ZNG1*, *Cre03.g175700*, *Cre12.g536900* and *Cre14.g613200*, were not affected by either. We also probed *CIA5* and *ZNG3* expression over the course of a day in phototrophically grown cultures that had been synchronized by 12h dark/12h light cycles [64]. We noted that *Cia5* and *ZNG3* are tightly co-expressed, with mRNA abundance of both genes peaking directly after the onset of light (Figure 2D). Both transcripts anticipated light and transcript abundance declined sharply after a brief peak at 0.5h after lights were turned on (Figure 2D). *Cre16.g692901* again had a very similar transcription profile to *ZNG3* and Cia5, while other CobW proteins showed a different response (Supplemental Figure 2). Among the 3 CobW proteins without the GTPase domain (CobW C only) transcripts encoding for LCI15 showed a very similar expression pattern to ZNG3, both others were also Cia5-dependently induced in low CO_2_ but unlike *ZNG3*, *Cre16.g692901* and *LCI15 we*re also induced in low Zn (Supplemental Figure 2).

**Figure 2.**
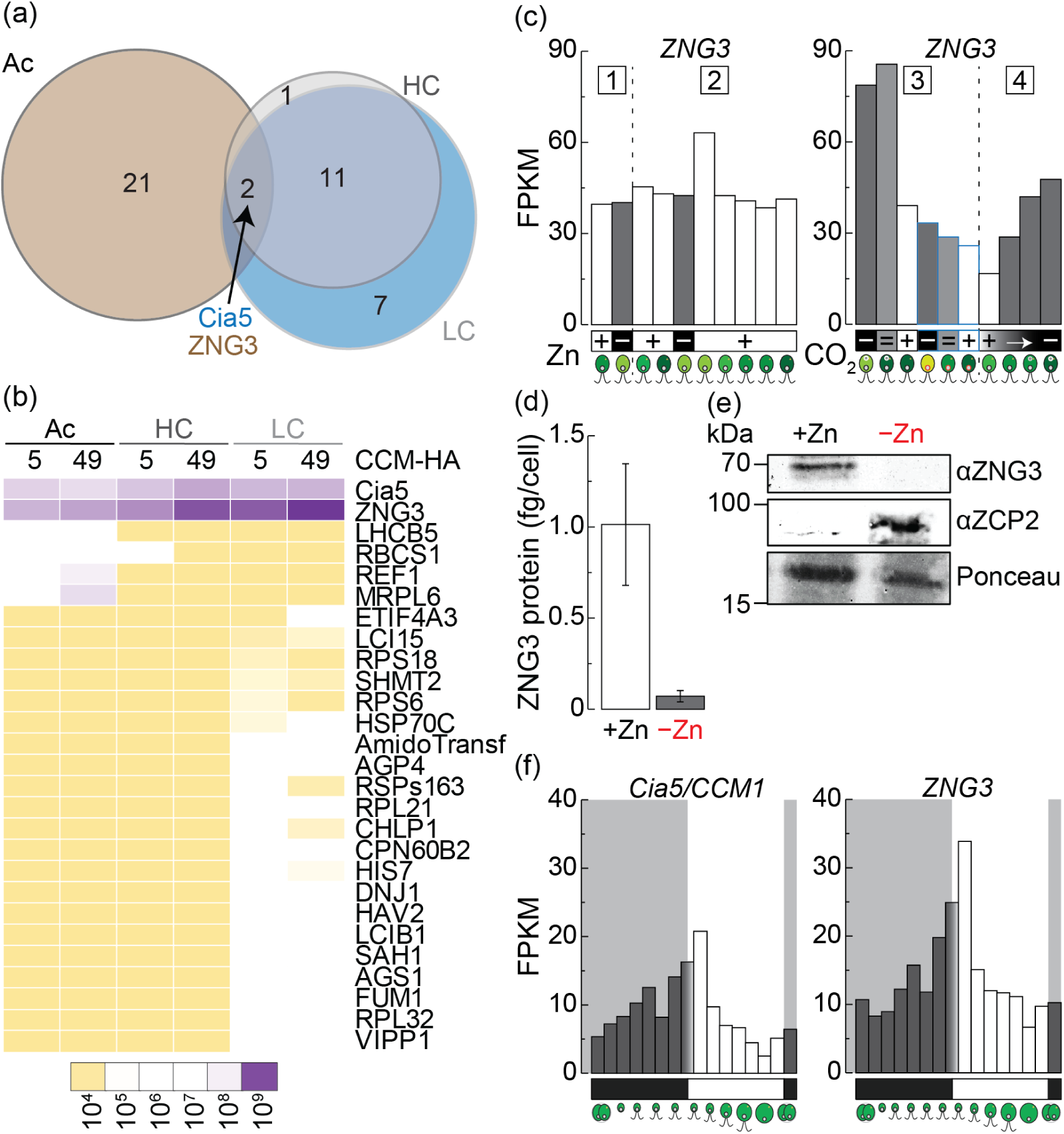
Cia5 constitutively interacts with ZNG3. (a) Immunoprecipitates of both Cia5-HA lines grown photoheterotrophically (Ac), or phototrophically (5% CO_2_, HC and 0.04% CO_2_, LC) were analyzed by LC-MS/MS. No. of identified proteins in both HA lines but not in wildtype controls are shown in circles. (b). Precursor ion intensities in Cia5-HA5 (5) and Cia5-HA49 (49) of proteins identified via LC-MS/MS (arbitrary units) in Ac, LC and HC conditions. (c) Survey of transcript abundances of *ZNG3* in published RNAseq datasets with varying Zn and CO_2_ supply. Malasarn *et al.* (1) analyzed transcripts in Zn-replete (+) and Zn-deficient (−) cultures, Hong Hermesdorf *et al.*, analyzed early exponential and early stationary Zn-replete cultures (+), as well as Zn-deficient (−) cultures and Zn resupply. Fang *et al.* (3) analyzed transcript abundances in cultures acclimated to high (+, 5%), air-level (=, 0.04%) and low CO_2_ (−, 0.01%) in wildtype (black outline) and *cia5* mutants (blue outline), while Brueggemann *et al.* analyzed the transition from high (+, 5%) to very low CO_2_ supply (−, 0.01%). (d) Absolute protein abundance of ZNG3 in mass spectrometry data from Zn-replete (+Zn) and Zn-deficient conditions (Strenkert *et al.*, 2023). (e) Total protein was extracted from wildtype (wt) in from Zn-replete (+Zn) and Zn-deficient (−Zn) conditions. Denatured proteins were separated with SDS-PAGE, transferred to nitrocellulose and immunodecorated with antibodies against ZNG3. Antibodies specific to ZCP2 were used as marker protein for Zn limitation, antibodies specific to CF_1_ served as loading control. (f) Survey of transcript abundances during the diurnal cycle (Strenkert *et al.* 2019).

Taken together, ZNG3 is a CobW-domain containing GTPase with a conserved metal-binding motif (GCxCC). Unlike *ZNG1* and *ZNG2*, transcript abundance of *ZNG3* is not affected by the Zn status of the cultures but solely responds to low CO_2_ stimulus, in a Cia5-dependent manner. While transcripts are unaffected by a lack of Zn, the protein is not present in low Zn environments. The Chlamydomonas genome encodes for two other CobW proteins with a similar expression profile, one with the GTPase domain present (Cre16.g692901) one with only the CobW C domain (LCI15).

### ZNG3 is important for growth when CO_2_ is not limited

To determine the involvement of the metal GTPase ZNG3 during the establishment of the CCM and in Zn homeostasis, we generated independent *zng3* mutants using CRISPR gene editing. Since our initial attempts to introduce *in-frame* stop codons into the *ZNG3* gene were not successful, we initially assumed that ZNG3 may be essential and aimed to generate inducible *zng3* mutants instead. To this end, we used a gRNA targeting the 5’UTR of *ZNG3* and an ssODN containing homology arms and a sequence encoding the *THI4* riboswitch. The native *THI4* riboswitch has been characterized before and represses gene expression of *THI4* when thiamine is present in the growth medium [65]. While the full-length incorporation of the *THI4* riboswitch into the 5’UTR of *ZNG3* was unsuccessful, we obtained two mutants, designated *zng3-1* and *zng3-2* which we selected for subsequent studies after genotyping. *zng3-1* harbors an insertion of a truncated, non-functional 442 bp *THI4* riboswitch fragment within its 5’UTR, containing several ORFs in all reading frames, while *zng3-2* suffered a 16 bp deletion that included the *ZNG3* start codon (Figure 3A). In immunodetections on total cell lysates of wildtype, *zng3-1* and *zng3-2* using a polyclonal ZNG3 antibody, *zng3* mutants appeared to be devoid of residual ZNG3 protein (Figure 3B). Like Cia5, ZNG3 was not differentially expressed in cells that were grown in LC, HC and Ac (Figure 3B). To test the effect of a loss of function in *zng3* on the ability of cells to grow in different carbon regimes, we grew *zng3* mutants alongside wildtype strains photoheterotrophically or phototrophically in HC and LC conditions (Figure 3CD). For comparison, we included the two independent *cia5* mutants in these experiments. We hypothesized that Zn delivery to Cia5 by ZNG3 would be required for successful CCM establishment and therefore expected *zng3* mutants to grow slower in low CO_2_ conditions, similar to *cia5* mutants as shown previously [27]. We recapitulated the expected growth phenotypes of *cia5* mutants, which were indeed asymptomatic when grown photoheterotrophically or phototrophically with high CO_2_ (5%) supply, but had reduced growth rates when CO_2_ became limiting (0.04%). However, *zng3* mutants did not phenocopy *cia5* mutants. Surprisingly, phototrophic growth of LC grown *zng3* mutants was not reduced but remained comparable to wildtype, with an average doubling time of 22h in *zng3* mutants as compared to 21h in wildtype cells. However, *zng3* mutants did exhibit a significant growth defect when grown phototrophically with high CO_2_ (HC) or when grown photoheterotrophically (Ac, Figure 3CD). This indicates that ZNG3 is not required for successful establishment of the CCM but does have a critical function in HC and in photoheterotrophic conditions, where Cia5 is not required. This also suggests that Cia5 is functional in strains lacking ZNG3, meaning Zn allocation towards Cia5 via the GTPase is not essential for successful induction of a CCM. We also analyzed the Zn content in *zng3* mutants in all growth regimes to determine if *zng3* mutants were still able to dynamically adjust their Zn quota in response to low CO_2_ availability, which was not the case. We found that differences in the intracellular Zn levels in *zng3* mutants relative to wildtype were not significant in these conditions but *zng3* mutants had higher levels of intracellular Zn in HC (Figure 3E).

**Figure 3.**
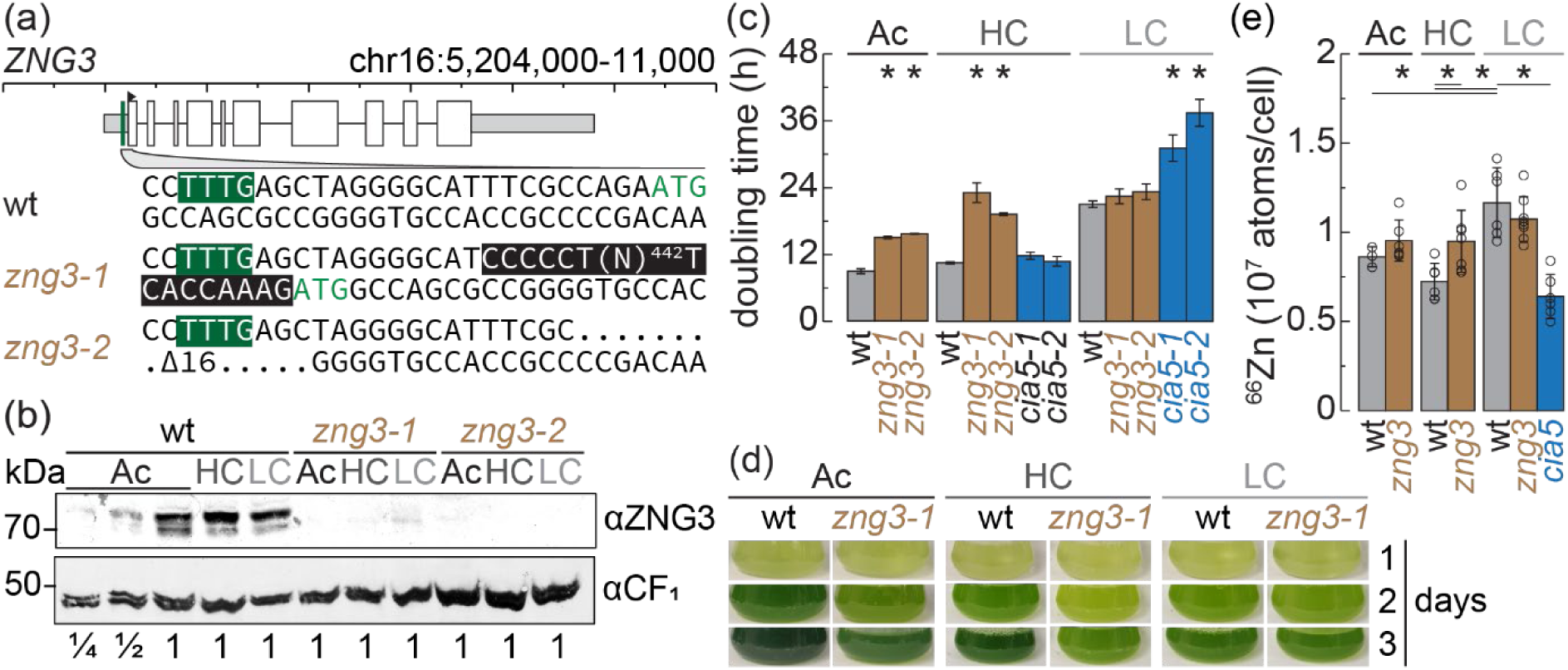
*zng3* mutants show reduced growth in high CO_2_ (5%). (a) Illustration of the *ZNG3* gene model showing the utilized PAM site (green) to generate CRISPR mutants and introduced gene edits in *zng3-1* and *zng3-2*. (b) Wildtype strains (wt), *zng3-1* and *zng3-2* mutants were grown photoheterotrophically (with acetate as carbon source, Ac), or phototrophically with supplementation of either 5% CO_2_ (HC) or 0.04% CO_2_ (LC). (b) Total protein extracts were separated by SDS PAGE and immunodetected using antisera against ZNG3. Antibodies specific to CF1 served as loading control. (c) Wildtype (wt), *zng3-1*, *zng3-2*, *cia5-1* and *cia5-2* were grown as in (b). Cell counts were obtained at 24-h intervals. Doubling times were calculated from data obtained during exponential growth. Shown are averages and standard deviation from three independent grown cultures. (d) shown are representative pictures of culture flasks taken on day 3 after inoculation. (e) Wildtype (wt) and *zng3* mutants were grown in HC, LC and photoheterotrophically (Ac, with acetate as a carbon source). Zn content was determined by ICP-MS/MS and normalized to cell numbers. Shown are averages and standard deviation of 6-9 independent experiments, as well as individual data points.

Taken together, ZNG3 mutants are similar in growth and Zn content to wildtype with no additional carbon supply, but show reduced growth when acetate or CO_2_ are provided.

### The plastid protein quality control pathway is induced in *zng3* mutants

To decipher the impact of loss of ZNG3 on Zn and carbon metabolism, we compared transcriptomes from wildtype cells and the two independent *zng3* mutants that were grown phototrophically with either air levels of CO_2_ (0.04%, LC), high CO_2_ (5% CO_2_, HC) or photoheterotrophically with acetate (Ac). We used the overlap of transcripts that were found either consistently reduced or induced in both mutants (log_2_ fold change > 1, FDR < 5%) to identify differentially expressed genes (DEGs) in Ac (481 up, 913 down), HC (149 up, 120 down) and LC (126 up, 165 down, Figure 4AB). Overall, between 0.6% and 5.2% of genes were differentially expressed between wildtype and *zng3* mutant strains with the largest fraction of DEGs found in photoheterotrophic grown cells (Figure 4A). We captured the expected transcriptional response when cells were grown in LC vs HC, including the characteristic induction of genes coding for the periplasmic carbonic anhydrase *CAH1*, HCO_3_ ^-^/CO_2_ importers *HLA3* and *LCI1*, chloroplast bicarbonate importer *LCIA*, [66] and mitochondrial CAs *CAH4/5* (Figure 5A and Supplemental Data 3). While all of the known low CO_2_ inducible (*LCI*) genes were still induced in *zng3* mutants in LC conditions, we observed minor changes in the extend of expression (Figure 5A). This included genes encoding the putative bicarbonate importer *HLA3* [67], the outward proton pump *ACA4* [68], *LCIC* [69] and the mitochondrial carbonic anhydrases *CAH4* and *CAH5* that have been shown to be necessary for photosynthetic growth in LC [54]. The expression of these genes is dependent on Cia5 [54], which suggests that Cia5 was still (partially) functional in *zng3* mutants, consistent with *zng3* mutants being asymptomatic when grown in low CO_2_ (Figure 3D). While in LC conditions the expression of almost all genes involved in the CCM was mildly to significantly reduced, some CCM-related genes, including *LCR1*, *LCIC*, *BST1* and *CCP2*, were induced in Ac, but the level of induction there did not reach the transcript amounts found in LC conditions (Figure 5A). We also noted that *EPYC1*, whose gene product is crucial for CCM establishment, was specifically downregulated in low CO_2_ grown *zng3* [19, 70] (Figure 5A).

**Figure 4.**
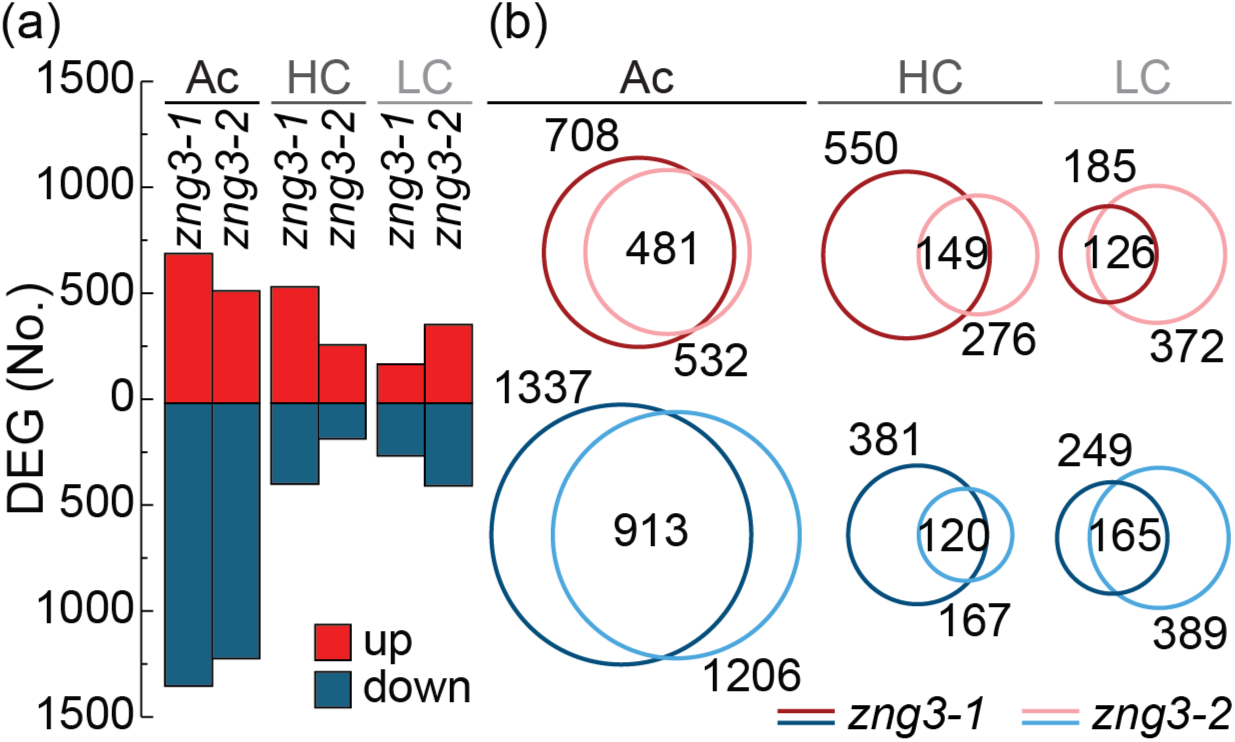
Differentially expressed genes between wildtype and *zng3* mutants. (a,b) Wildtype strains (wt), *zng3-1* and *zng3-2* mutants were grown photoheterotrophically (with Acetate as the carbon Source, Ac), or phototrophic with supplementation of either 5% CO_2_ (HC) or 0.04% CO_2_ (LC). (a) Bar graph showing number of differentially expressed genes between *zng3-1* or *zng3-2* and wildtype. (b) Venn diagrams identify the number of differentially accumulating transcripts at ≥2-fold change (FDR < 0.05) in each pairwise comparison and the respective overlap of different sets.

**Figure 5.**
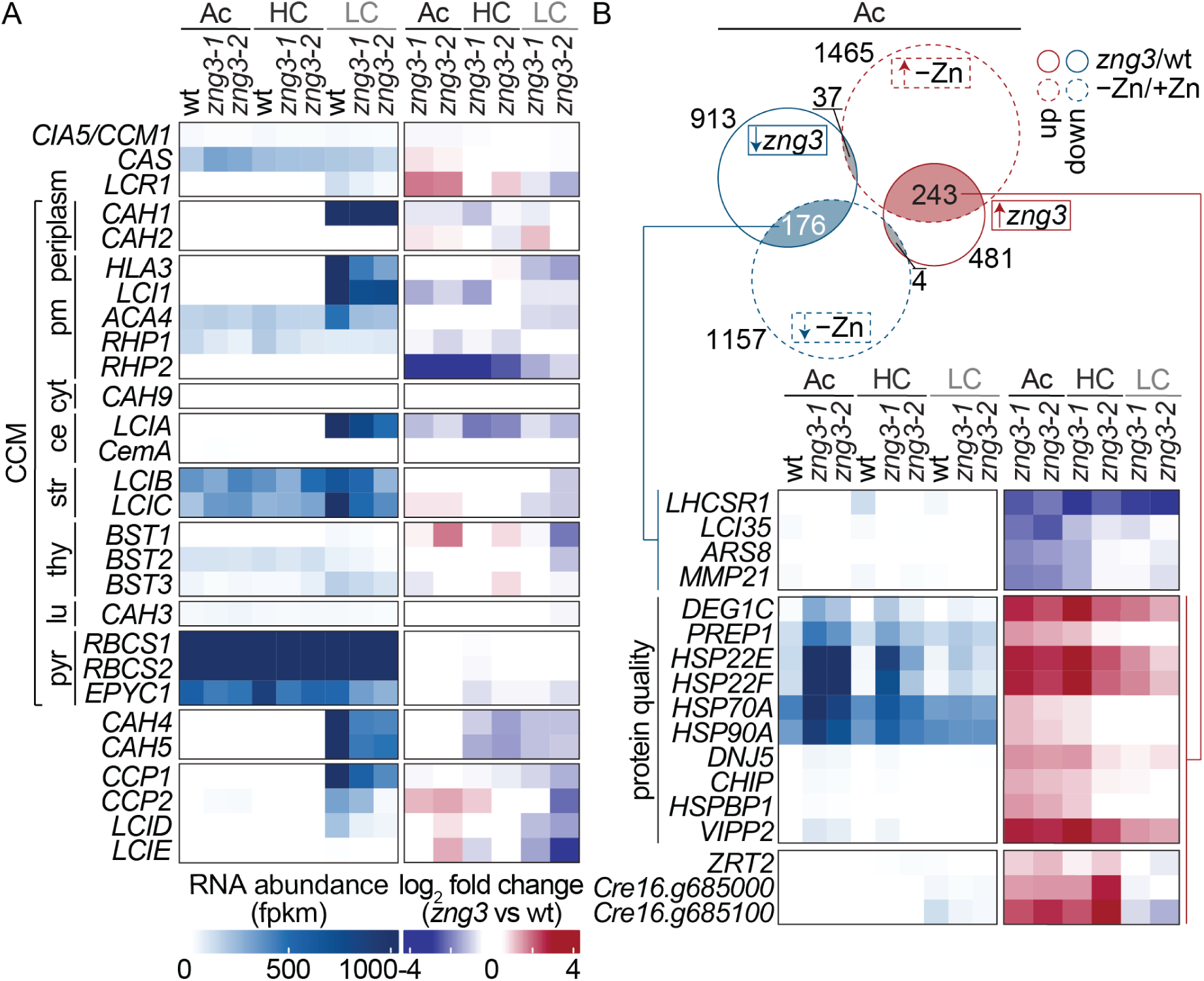
Genes encoding proteins involved in plastid quality control and Zn metabolism are induced *zng3* mutants. (a) Shown are amounts and changes to transcript abundances (RNAseq) of genes encoding for proteins involved in the CCM. The left heatmap illustrates transcript abundances (fpkm), the heatmap on the right shows the log_2_ fold changes between *zng3* mutants and wt, cultured photoheterotrophically (acetate, Ac) or phototrophically bubbled with 5% CO_2_ (HC) or air (LC). Shown are averages of 6 wildtype cultures (2 independent strains, 3 replicates each) and 3 independent cultures of each mutant. (b) Overlap of differentially accumulating transcripts (RNAseq) between Zn deficient and Zn replete cultures (from Hong-Hermesdorf *et al.* 2014) and between *zng3* mutants and wildtype cultures (this study). Heatmaps and samples follow the scheme described in (a). Examples shown are genes with a consistent, significant response in both *zng3* mutants in at least one growth mode. A more expansive list of genes involved in these pathways, including those not changing in *zng3* mutants, can be found in Supplemental Figures 4-6.

Since *ZNG3* encodes a candidate Zn handling GTPase, we expected expression of genes coding for proteins involved in Zn homeostasis to be potentially impacted in *zng3* mutants. Indeed, we observed a specific and significant transcriptional upregulation of two genes encoding unannotated CobW-C domain containing proteins (Cre16.g685000 and Cre16.g685100, Supplemental Figure 1) in HC and Ac grown *zng3* mutants as compared to wildtype (Figure 5A). Transcripts encoding for both proteins were Zn-dependently induced [47], and increased transcript abundance in low CO_2_ grown cells was dependent on Cia5 (Supplemental Figure 3). Both genes are located close to each other on chromosome 16, interspersed only by another CobW-C domain containing protein (*LCI15*) that is not responsive to low Zn but also was Cia5-dependently increased in LC (Supplemental Figure 3). This data suggests that a loss of ZNG3 function subsequently impacts other, specific Zn handling components that may be upregulated to compensate for a loss in ZNG3 function. Expression of other genes encoding proteins involved in Zn import and distribution, including ZRT1/2/3, ZCP2, ZNG2, Cre12.g536900 and Cre16.g658000 were not induced as much as the two other CobW-C containing proteins in *zng3* mutants, but did show some increased expression in Ac and/or HC conditions (Figure 5B, Supplemental Figure 4). While significant, the increased *ZRT1*, *ZRT2* and *ZRT3* expression in *zng3* mutants was amplitudes lower as compared to mRNA abundance of these genes in Zn limited grown cells [47] (Supplemental Figure 3).

Photoheterotrophically grown *zng3* mutants are most severely impaired in growth and transcript accumulation as compared to wildtype cells (Figure 4A). We therefore compared the transcriptional response in these strains with the response to Zn-limitation in a previous study [47]. We found that a substantial fraction of the differentially accumulating transcripts in Zn limited cells was also differentially accumulated in Zn-replete *zng3* mutants (Figure 5B). Transcripts accumulating in Zn-limitation also accumulated in *zng3* mutants and *vice versa*. Genes coding for selected chaperones involved in plastid protein quality control were amongst the genes accumulating in Zn deficient cells and *zng3* mutants (Supplemental Figure 6). Transcripts of chloroplast small heat shock proteins (sHSPs) HSP22E and HSP22F were dramatically up-regulated in *zng3* mutants, with the amplitude of induction being higher in conditions in which *zng3* mutants show a significant growth defect (Ac and HC, Figure 4A, Supplemental Figure 6). Heat shock proteins belonging to that family bind to unfolded or partially folded proteins, to potentially facilitate either refolding or at least to prevent aggregate formation followed by targeted degradation. HSP22E/F chaperone activity is crucial for maintaining plastid protein homeostasis during heat stress in Chlamydomonas, but identified clients of the sHSP include proteins involved in starch biosynthesis and CO_2_ assimilation [71]. The increase in transcripts encoding for chloroplast targeted sHSPs could imply that *zng3* mutants accumulate unfolded proteins within the chloroplast. In support of this hypothesis, a chloroplast localized protease, DEG1C [72], which is involved in plastid protein quality control, was also up-regulated in *zng3* mutants, with the amplitude of induction again being higher in Ac and HC grown cells. Interestingly, it has been shown previously that a small subset of proteins involved in establishing the CCM, like plastid LCIB and LCIC but also LHCSR1 accumulate in *deg1c* mutants, indicating their dependence on DEG1C for quality control [72]. Transcripts encoding for LCIC (only in LC) and LHCSR1 (all conditions) were both found to be reduced in *zng3* mutants.

In summary, our transcriptome analysis suggests that, with few exceptions, components of the CCM are generally not misregulated in *zng3* mutants in LC conditions, indicating that ZNG3 is not required to establish a CCM in Chlamydomonas. Instead we found a substantial overlap of genes induced in *zng3* mutants to Zn deficiency, indicating that ZNG3 is involved in coordinating a response to low Zn environments. Most prominently we found two metallochaperones that, like ZNG3, contain a CobW-C domain, highly induced in *zng3* mutants, indicating that intracellular Zn trafficking is adjusted ZNG3-dependently. Both metallochaperones are Cia5-dependently induced in low CO2 environments, the derepressing of these genes in *zng3* mutants points to a role of ZNG3 in preventing their expression in Zn replete conditions. Chloroplast-targeted small heat shock proteins and a chloroplast Deg protease were found highly induced in *zng3* mutants grown photoheterotrophically and with additional CO_2_ supply. These proteins have been implicated for roles in the CCM through some of their own target proteins. While this could be an indirect outcome to prevent accumulation of misfolded or mis-metalated proteins in Ac and HC conditions as a consequence of ectopic metallochaperone expression, this could also be part of a direct, ZNG3-coordinated effort to in tailoring the CCM with regards to Zn demand. Taken together, the physical connection between ZNG3 and CIA5 offers a platform to coordinate Zn and CO_2_ acclimation.

## DISCUSSION

Since Zn is a crucial cofactor for many proteins, organisms have evolved sophisticated systems that include Zn uptake, intracellular distribution, compartmentalization and efflux to achieve Zn homeostasis [73]. In recent years, we have started to elucidate the fascinating components responsible for Zn transport and sequestration, but mechanisms for Zn delivery to target proteins, while implied by the existence of CobW proteins as hypothetical Zn chaperones, still remain largely unknown [49, 73].

Despite the striking evolutionary conservation, the functions of CobW proteins other than for the vertebrate, yeast and plant ZNG1 [50, 51], remain completely unknown because we have limited information regarding potential client proteins. In this work, we have identified Cia5 as a direct ZNG3 target using immunochemistry and mass spectrometry-based protein identification. Based on previous work on the vertebrate and yeast ZNG1, our assumption is that ZNG3 interacts with the Zn finger domain of Cia5 and uses GTP hydrolysis to transfer Zn to the second potential Zn binding site. It was proposed that GTP hydrolysis drives Zn transfer from ZNG1 to the catalytic site of its client, the Zn metalloprotease METAP1. As a result, METAP1 activity is restored by ZNG1 if cellular Zn is low [51]. We do not see differential association between Cia5 and ZNG3 when cells are grown under different carbon regimes, implying that metalation of Cia5 by ZNG3 does not necessarily result in dissociation between the Zn chaperone and its client after GTP hydrolysis but rather may have other, structural implications. In addition, mutants in *zng3* do not phenocopy *cia5* mutants, grow like wildtype in low CO_2_ environments and, with few exceptions, upregulate expression of low CO_2_ induced genes just like wildtype, suggesting that Cia5 activity does not rely on ZNG3 function to induce establishment of the CCM. We propose that ZNG3 plays a role in adjusting Zn homeostasis in response to a reduced Zn demand when Chlamydomonas does not need a CCM. This hypothesis is consistent with the observation that expression of genes encoding chaperones involved in plastid quality control is upregulated in *zng3* mutants that are grown in conditions in which the CCM is not induced, suggesting protein unfolding and aggregation within this organelle. We assume that Zn might be mistakenly sequestered to the plastid in *zng3* mutants even when a CCM is not formed and plastid Zn demand is low, resulting in mis-metalation of non-Zn proteins and protein mis-folding. Our data suggests that the Zn handling GTPase ZNG3 represents a crucial, regulatory off switch to adjustments to Zn metabolism that are needed during establishment of the CCM.

The discovery of the nuclear ZNG3-Cia5 complex and the results we have presented offer novel insights into the role of CobW proteins in regulating carbon metabolism in general and potential avenues of Cia5 activation specifically. But we need additional biochemical and structural characterizations to understand how this unusual, regulatory ZNG3-Cia5 pair balances Zn and carbon metabolism.

## ACKNOWLEDGMENTS

We thank the staff at the MSU Genomics Core for help with RNA quality control, cDNA sample preparation, library preparation and sequencing. We are thankful to the MSU proteomics core facility and Douglas Whitten for mass-spectrometry based protein identification of immunoprecipitated samples. This research was partially supported by a fellowship from Michigan State University under the Training Program in Plant Biotechnology for Health and Sustainability (T32-GM152798 and T32-GM110523). This work was also supported by grants from the National Institutes of Health for the resource entitled “Quantitative Elemental Mapping for the Life Sciences” (P41 GM135018) and from the U.S. Department of Energy, Office of Science, Basic Energy Sciences (DE-FG02-91ER20021).

## AUTHOR CONTRIBUTIONS

D.S. and St.S. designed the research; G.K.A., St.S., Sa.S., A.M. performed research; G.K.A., St.S., T.V.O and D.S. analyzed and interpreted data; D.S. and S.R.S. wrote the paper all authors edited and approved the final version of the manuscript.

## MATERIAL AND METHODS

### Chlamydomonas growth conditions

Chlamydomonas parental strain used for all experiments was CC-425. Cells were grown in tris-acetate-phosphate (TAP) medium or high salt medium from Sueka (HSM) and prepared with revised trace elements [74] with constant agitation at 120 rpm in a shaker incubator (Multitron, Infros HT, Annapolis Junction, MD, USA) at 24°C in continuous light (60 µmol m^−2^ s^−1^) [74]. Phototrophically grown cultures were directly bubbled with humidified air/CO_2_ mixtures (5% CO_2_, HC or 0.04% CO_2_, LC) as indicated.

### Generation of Chlamydomonas mutant strains using CRISPR/Cpf1

Chlamydomonas CC-425, a cell-wall reduced arginine auxotrophic strain, was used for transformation with a *Hin*DIII digested pMS666 containing the *ARG7* gene and a ribonucleoprotein (RNP) complex consisting of a guide RNA (gRNA) targeting a protospacer adjacent motif (PAM, TTTV) sequence and LbCpf1 as described in [52, 53, 75]. ssODN templates with the intended edits were provided prior to electroporation. Wildtype control strains used in this study were CC-425 transformed alongside mutants providing a *Hin*DIII digested pMS666 containing the *ARG7* gene only to restore arginine prototrophy. PAM sequences and gRNAs are listed in Supplemental Table 1. Transformants selected on arginine-free TAP plates were screened via colony PCR. For this purpose, Chlamydomonas cells were transferred from TAP agar plates and resuspended in 100 μL of TE buffer (10 mM Tris, 1 mM EDTA, pH 8.0). The cell suspension was then heated at 95°C for 10 minutes followed by vigorous vortexing and centrifugation at maximum speed for 5 minutes. The supernatant was used for Polymerase Chain Reaction (PCR) for genotyping using either GoTaq Green Mastermix (Promega, Medison, WI, USA) with the addition of 1 M Betaine or for quantitative PCR (qPCR) with SYBR Green Mastermix (Bio-Rad) and addition of 1 M Betaine. The primer sequences used for screening the transformants are also provided in Supplemental Table 1. Screening for successful integration was done using diagnostic primers that specifically amplify either transgenic or endogenous DNA sequences at the integration site. For screening of Cia5-HA strains, individual colonies were grown on TAP media and total cell lysates were separated on SDS-PAGE, transferred to nitrocellulose membranes and decorated using anti-HA antibody as described below.

After selecting candidates (mutant strains or strains expressing tagged proteins, DNA sequences were assessed by amplification of the gene edited region with sequencing primers (Supplemental Table 1) and using GoTaq Green Mastermix (Promega, Medison, WI, USA) with the addition of 1 M Betaine. The PCR products were loaded into 1% agarose gel for gel electrophoresis with TAE buffer. The DNA fragment was then extracted using the E.Z.N.A. Gel Extraction Kit (Omega Bio-tek) according to the instructions and analyzed by sanger sequencing.

### Quantitative elemental analysis

Cell cultures, during exponential growth at a culture density of 3–8 × 10^6^ cells/mL, corresponding to 5 × 10^7^ cells were collected by centrifugation at 1424*g* for 3 min in a 50-mL falcon tube. The cells were washed once in 50 mL of 1 mM Na_2_EDTA pH 8.0 (to remove cell surface–associated metals) and once in Milli-Q water. The cell pellet was stored at −20°C before being overlaid with 286 µL 70% (v/v) nitric acid and digested at 65°C over night before being diluted to a final nitric acid concentration of 2% (v/v) with Milli-Q water. Metal, sulfur and phosphorus contents were determined by inductively coupled plasma mass spectrometry (ICP-MS/MS) on an Agilent 8800 Triple Quadropole ICP-MS instrument by comparison to a calibration standard (Agilent 5183-4688). ^45^Sc served as an internal standard (Inorganic Ventures MSY-100PPM). The levels of ^66^Zn were determined in MS mode directly using He in a collision reaction cell. An average of three technical replicate measurements was used for each individual biological sample. The variation between technical replicate measurements never exceeded 5% for an individual sample. 3 to 9 samples from independent cultures in each individual conditions were used to determine the average abundance and variation between cultures shown in all figures; individual points indicate the abundance of each individual independent culture.

### Antibody Production and Immunoblotting

Antibodies targeting ZNG3 were produced by Labcorp by immunization of rabbits using the subcutaneous implant procedure of rabbits following a 118-day protocol with synthetic peptide Ac-ERNPKRRAKRLHDLC-NH_2_. The antibodies to ZNG3 were affinity purified by Labcorp using immobilized peptide. For immunoblot analysis, a total of 2x10^7^ Chlamydomonas cells were collected from cultures at mid-logarithmic phase by centrifugation at 3,000*g* for 3 minutes at 4°C. Protein quantification, the cell pellets were resuspended in 50-200 µL of 10 mM sodium phosphate buffer pH 7.0 (4.23 mM NaH_2_PO_4_, and 5.77 mM Na_2_HPO_4_) with or without protease inhibitor cocktail (Roche) and kept at -80°C until further use. Total protein concentration was determined using the Bicinchoninic Acid (BCA) assay (Pierce™ BCA Protein Assay Kits, Thermo Scientific, Waltham, MA, USA). Samples were mixed with 1 volume of 2x Laemmli’s sample buffer 125 mM Tris-HCl pH 6.8, 20% Glycerol, 4% SDS, 10% β-Mercaptoethanol, and 0.005% Bromophenol blue) and incubated at 65°C for 20 minutes, followed by ice incubation for 2 minutes. Proteins were separated by denaturing PAGE and transferred to nitrocellulose. Unless noted otherwise, 10 μg total protein was loaded on each acrylamide gel. The protein was separated using SDS-PAGE and blotted onto 0.4 μm nitrocellulose membrane for western blotting using a Power Blotter (Invitrogen). The membranes were incubated with 3% non-fat dried milk (w/v) in PBST (137 mM NaCl, 2.7 mM KCl, 10 mM Na_2_HPO_4,_ 2 mM K_2_HPO_4_, and 1% Tween-20) for 1 h. The membranes were then incubated in the primary antibody: anti-HA (1:10000), anti-ZNG3 (1:1000), anti-ZCP2 (15000, Agrisera AS12 1848), anti CF_1_ (1:40000, gift from Sabeeha Merchant) in 3% non-fat dried milk (w/v) in PBST with constant agitation. The membranes were then washed 3 times with PBST and subjected to incubation with a goat anti-rabbit antibody conjugated with alkaline phosphatase at 1:8000 (v/v) in 3% non-fat dried milk (w/v) PBST for 1 h with constant agitation. The secondary antibody used was a goat anti-rabbit antibody conjugated to alkaline phosphatase and processed according to the manufacturer’s instructions.

### RNA extraction and qRT-PCR

A total of 5-8x10^7^ cells were collected by centrifugation at 1610 *g* for 2 minutes at 4°C. The cell pellet was resuspended in 1 mL of TRIzol^®^ Reagent (Invitrogen). Two hundreds μL of chloroform were added, followed by mixing and centrifugation at 13,200 rpm (Eppendorf, 5430 R) for 15 minutes at 4°C. Five hundred μL of the supernatant was transferred to a new tube containing 700 μL of isopropanol. The mixture was incubated on ice for 15 minutes followed by centrifuging at 12,000 rpm (Eppendorf, 5430 R) for 15 minutes at 4°C. The resulting pellet was dried and resuspended in 40 μL of RNase-free water. DNA digestion and RNA clean-up were carried out with the Zymo Clean & Concentrator 5 kit (Zymo Research) by following the instruction manual. cDNA was synthesized with M-MLV Reverse Transcriptase (Invitrogen). qRT-PCR was performed using SYBR Green MasterMix. *RACK1* served as a loading control. Sequences of the *CAH4* qRT-PCR sequences are in Supplemental Table 1. The PCR cycle is as followed; 95°C for 10 minutes, followed by 40 cycles of 95°C for 15 seconds and 65°C for 1 minute, 95°C for 15 seconds, 60°C for 1 minute, the temperature is then increased to 95°C at 1%/second, temperature is then reduced to 60°C and held for 1 minute, and reduced to 4°C for storage.

### RNA sequencing

Two independent wildtype controls and both *zng3* mutants were grown phototrophically (HSM medium) with either air levels of CO_2_ (0.04%, LC), high CO_2_ (5% CO_2_, HC) or photoheterotrophically (TAP medium) in the presence of the reduced carbon source acetate (Ac), each in triplicate independent cultures for each growth condition (36 samples). RNA was extracted as described above, abundance was determined using the Qubit RNA BR Assay Kit. Libraries were prepared using the Watchmaker Genomics mRNA Library Preparation Kit with IDT xGEN 10nt Unique Dual-Index primers following manufacturer’s recommendations.

Completed libraries were QC’d and quantified using a combination of Biotium AccuGreen High Sensitivity dsDNA and Agilent 4200 TapeStation HS DNA1000 assays. Libraries were normalized to a consistent concentration and equal amounts were pooled. The pool was quantified using the Invitrogen Collibri Quantification qPCR kit. Sequencing was performed using an Element AVITI Cloudbreak Freestyle High Output 300 cycle kit, in a 2x150bp paired end format. Base calling was done by AVITIOS v3.3.2 and the output was demultiplexed and converted to FastQ format using Element Biosciences bases2fastq v2.1.0.

Reads were aligned to the most recent version of the Chlamydomonas genome (v6) [58] using STAR (2.7.11b) [76]. Expression estimates in fragments per kilobase of transcript per million fragments mapped (FPKM) were determined by Cufflinks and differential expression analysis was performed with Cuffdiff (v2.2.1) [77]. Genes were considered significantly changing requiring a log_2_ fold change of at least ±1 and an FDR-adjusted p-value < 0.05. Subsequent data analysis and figure production were carried out in R, and Adobe Illustrator was used for further editing.

### Immunoprecipitation followed by Mass spectrometry (IP-MS/MS)

Approximately 1x10^9^ cells were collected by centrifugation at 1610 *g* for 3 min at 4°C. Cell pellets were washed twice with 15 mL of KH buffer (20 mM Hepes-KOH pH 7.2 and 80 mM KCl) and were resuspended in 1 mL of lysis buffer (20 mM Hepes-KOH pH 7.2, 1 mM MgCl_2_, 1 mM KCl, 15 mM NaCl, 1x protease inhibitor cocktail (Roche), and 0.1% triton X-100). Cells were broken by sonication on ice. Cell debris were removed by centrifugation at 1610 *g* for 3 min minutes at 4°C. The supernatant was transferred to a new 1.5 mL centrifuge tube and combined with Protein A Sepharose beads coupled with anti-HA antibodies. The protein-bead mixture was incubated with constant rotation at 4°C for 2 hours. The beads were then recovered by centrifugation at 5000 rpm for 20 s at 4°C, washed three times with lysis buffer (20 mM Hepes-KOH pH 7.2, 1 mM MgCl_2_, 1mM KCL, 15mM NaCl, 1x Protease Inhibitor Cocktail, 0.1% Triton) and twice with 10 mM Tris-HCl (pH 7.6). The supernatant was removed, and the beads were frozen for subsequent protein identification by mass spectrometry (MS). For proteolytic digestion, antibody-bound proteins were digested on-bead by washing 3 times using 50mM ammonium bicarbonate. Trypsin, in the same buffer, was then added to the beads at 5ng/uL so that the beads were just submerged in digestion buffer and allowed to incubate at 37°C for 6 h.

The solution was acidified to 1% with trifluoroacetic acid and centrifuged at 14,000 *g*. Peptide supernatant was removed and concentrated by solid phase extraction using StageTips [78]. Purified peptides eluates were dried by vacuum centrifugation and frozen at -20 °C or re-suspended in 2% acetonitrile/0.1%TFA to 20 µl. For LC/MS/MS analysis, an injection of 10 µl was automatically made using a Thermo (www.thermo.com) EASYnLC 1200 onto a Thermo Acclaim PepMap RSLC 0.1mm x 20mm C18 trapping column and washed for ∼5min with buffer A. Bound peptides were then eluted over 35min onto a Thermo Acclaim PepMap RSLC 0.075mm x 250mm resolving column at a constant flow rate of 300nl/min with a gradient of 8%B to 22%B from 0min to 19min and 22%B to 40%B from 19min to 24min. After the gradient the column was washed with 90%B for the duration of the run (Buffer A = 99.9% Water/0.1% Formic Acid, Buffer B = 80% Acetonitrile/0.1% Formic Acid/19.9% Water. Column temperature was maintained at a constant temperature of 50°C using an integrated column oven (PRSO-V2, Sonation GmbH, Biberach, Germany). Eluted peptides were sprayed into a ThermoScientific Q-Exactive HF-X mass spectrometer (www.thermo.com) using a FlexSpray spray ion source. Survey scans were taken in the Orbi trap (60000 resolution, determined at m/z 200) and the top 15 ions in each survey scan are then subjected to automatic higher energy collision induced dissociation (HCD) with fragment spectra acquired at a resolution of 7500. The resulting MS/MS spectra are converted to peak lists using Mascot Distiller, v2.8.5 (www.matrixscience.com) and searched against a protein sequence database containing Chlamydomonas entries provided by the customer appended with common laboratory contaminants (downloaded from www.thegpm.org, cRAP project) using the Mascot searching algorithm, v 2.8.3. [79] The Mascot output was then analyzed using Scaffold, v5.3.3 (www.proteomesoftware.com) to probabilistically validate protein identifications. Assignments validated using the Scaffold 1%FDR confidence filter are considered true. Mascot parameters for all databases were as follows: allow up to 2 missed tryptic sites, variable modification of Oxidation of Methionine, peptide tolerance of +/-10ppm, MS/MS tolerance of 0.02 Da, FDR calculated using randomized database search.

### Statistical analysis

Unless stated otherwise, a two-sided T-test was used to determine statistical differences between samples. Asterisks between samples indicate a p-value of < 0.05.

---

**Supplemental table 1.**
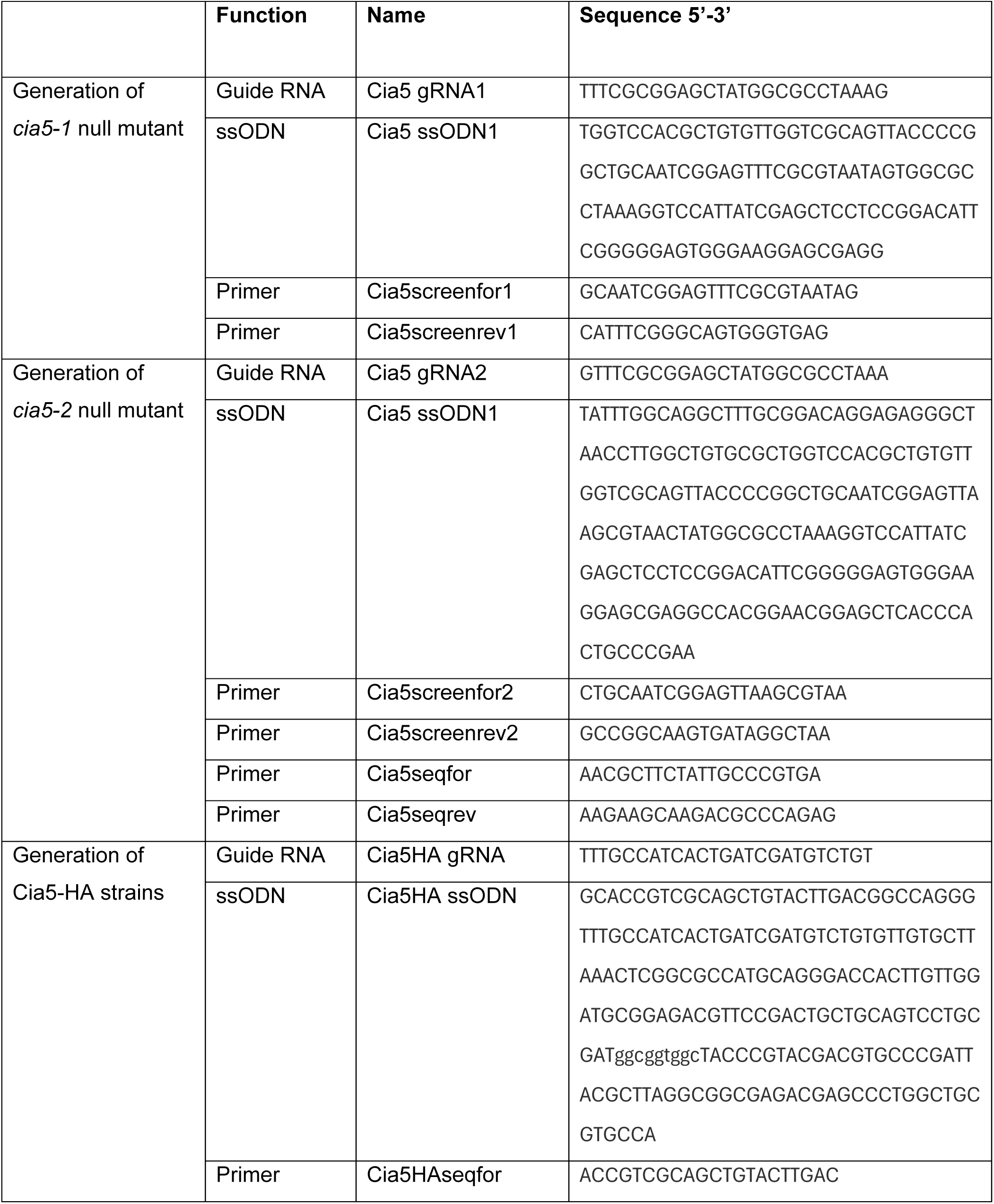
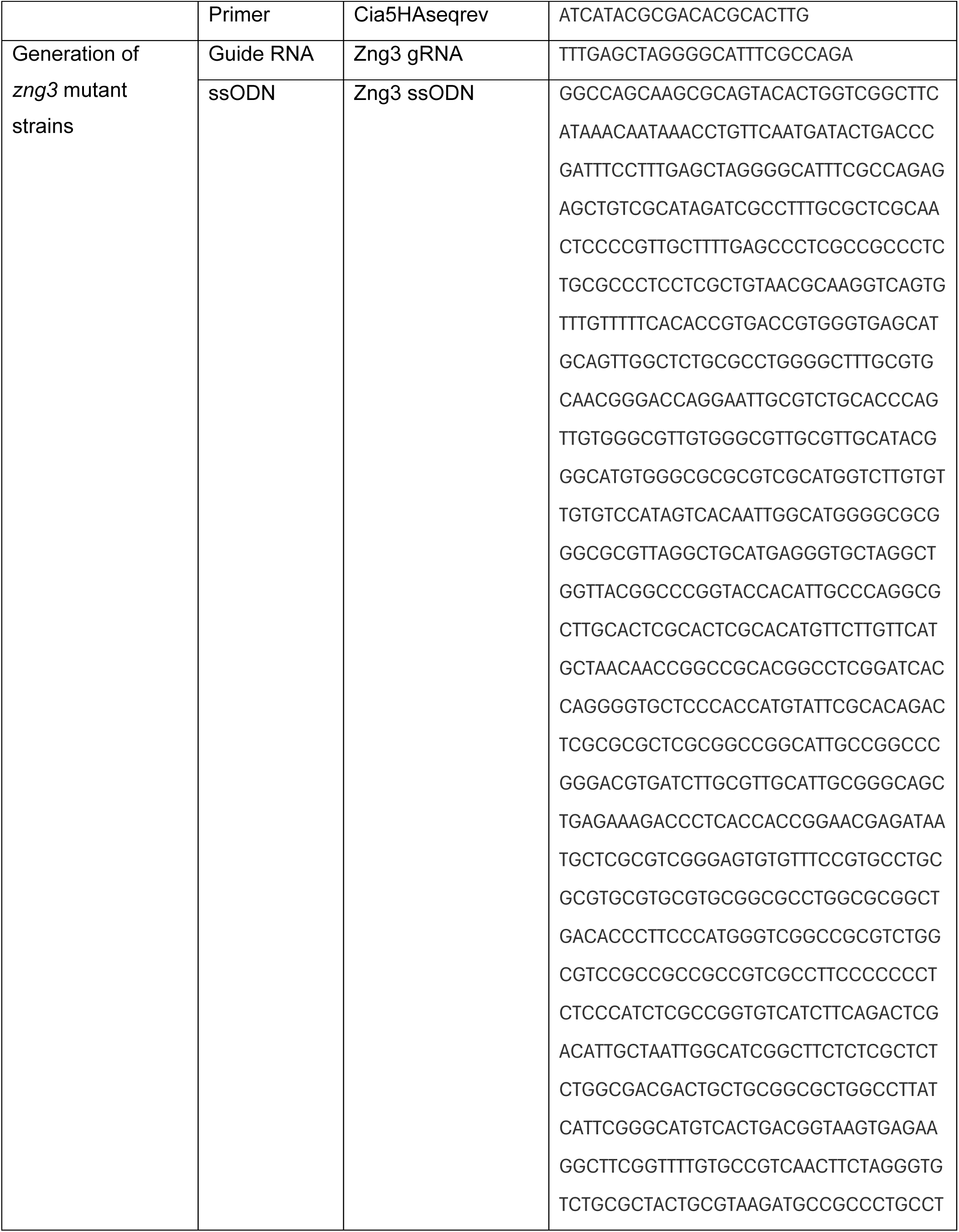
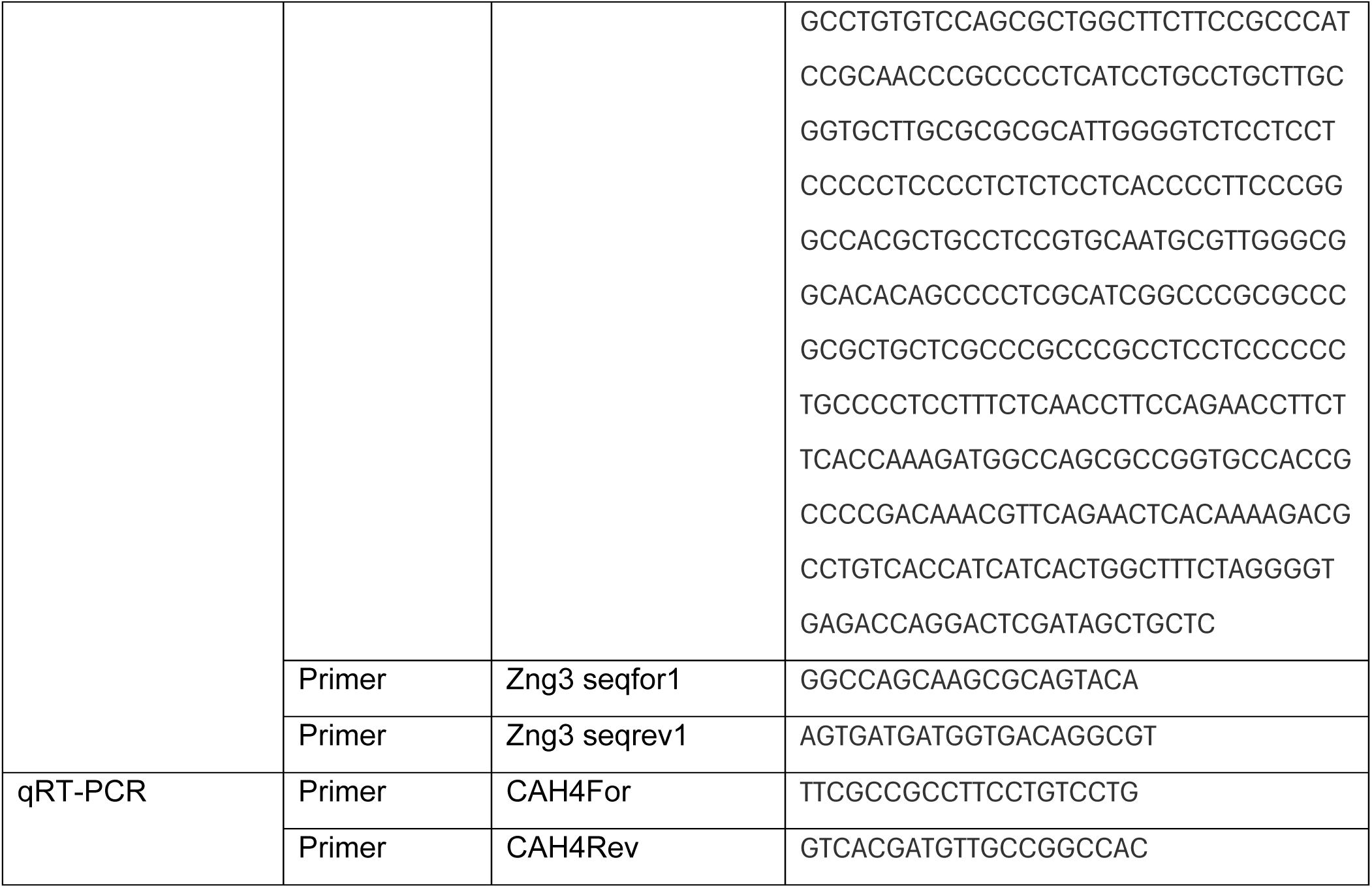
Sequences of gRNAs, primers and ssODNs used in this study.

