## Supplemental Figures for "CIA5 INTERACTS WITH THE ZINC CHAPERONE ZNG3 TO BALANCE CARBON AND ZINC METABOLISM"

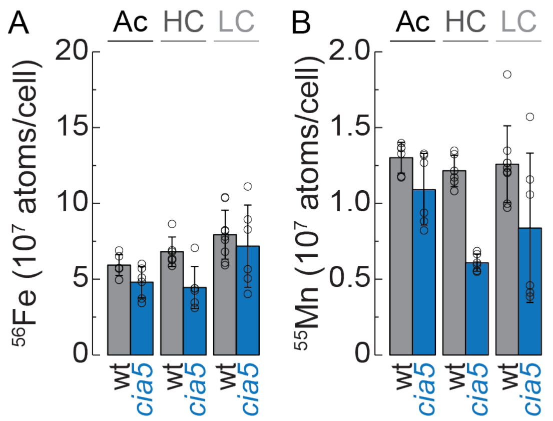


**Supplemental Figure 1. Fe and Mn content is unchanged in LC as compared to HC and Ac grown wildtype cells.** Wildtype (wt) and *cia5* mutants were grown in HC, LC and photoheterotrophically (Ac, with acetate as a carbon source). Fe and Mn content was determined by ICP-MS/MS and normalized to cell numbers. Shown are averages and standard deviation of 6 independent experiments, as well as individual data points.

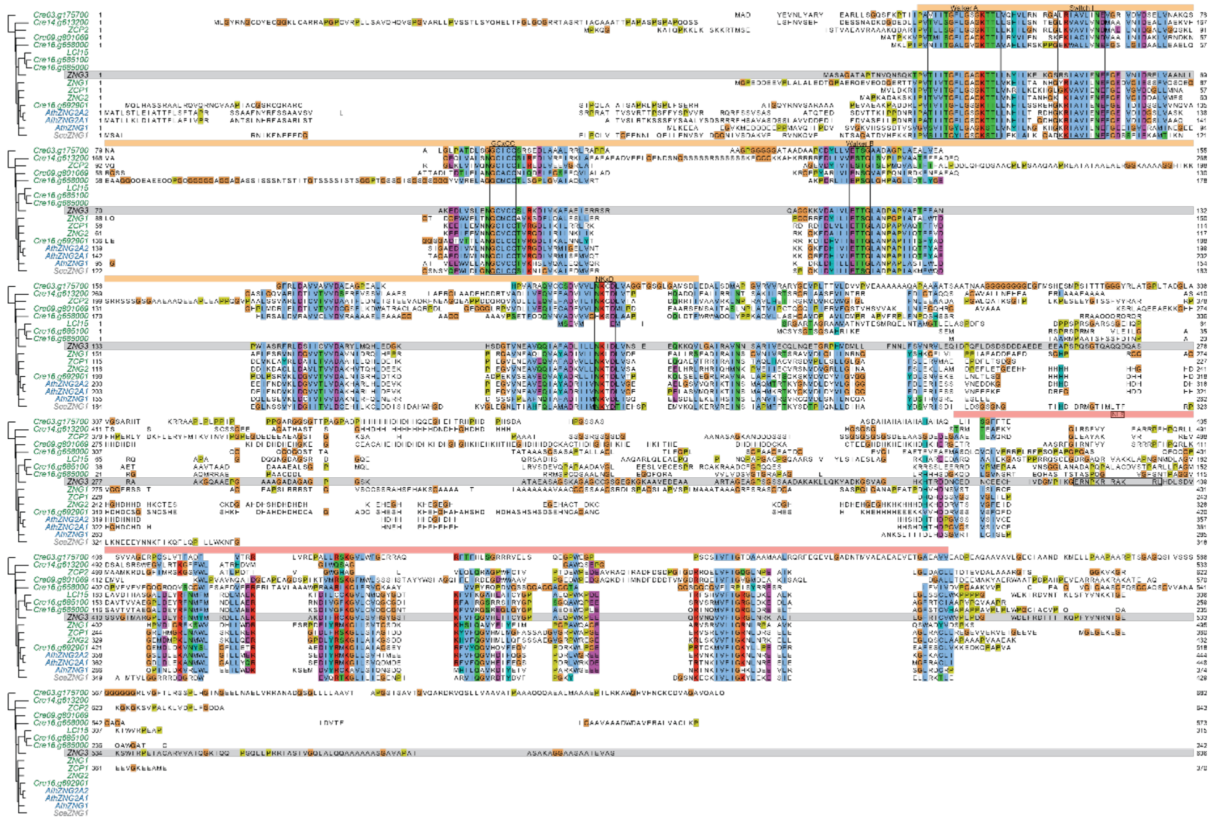

**Supplemental Figure 2. Multiple sequence alignment between CobW (COG0532) domain proteins in *Chlamydomonas reinhardtii*, *Arabidopsis thaliana* and *Saccharomyces cerevisiae*.** Sequences were aligned using Clustal Omega (https://pubmed.ncbi.nlm.nih.gov/21988835/) and organized in Jalview (https://doi.org/10.1093/bioinformatics/btp033). The CobW domain (N-terminal, GTPase) is indicated by an orange box above the protein sequences, the C-terminal CobW_C domain is indicated by a red box above the sequences. Important motifs are highlighted by a black outline and labelled accordingly. Chlamydomonas CobW proteins are labelled green, with the exception of ZNG3 , which is labelled black and its amino acid sequence in all paragraphs is highlighted by a grey background. Arabidopsis proteins are labelled in blue, *Saccharomyces cerevisiae* ZNG1 is labelled in grey.

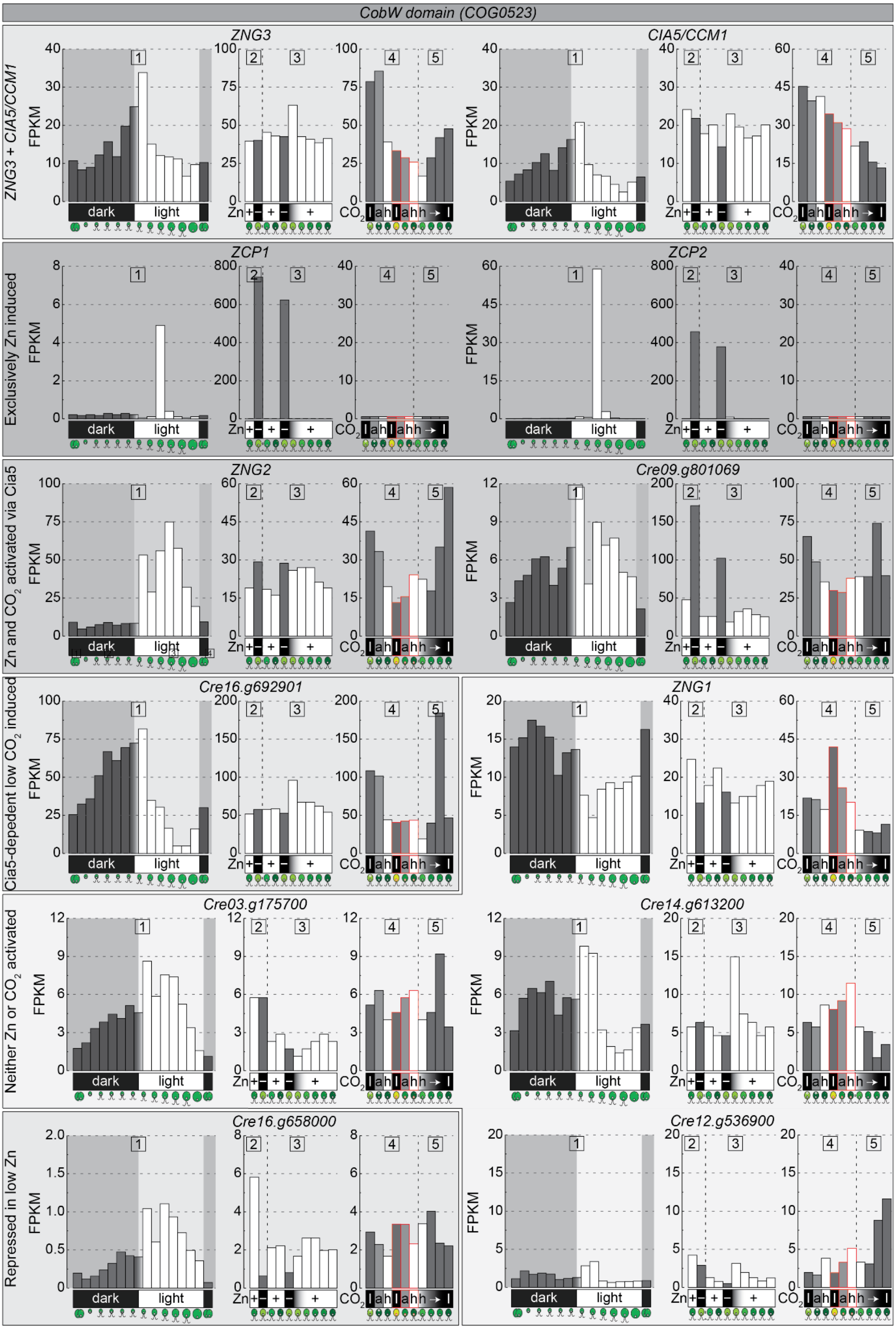
CobW domain (COG0523)
ZNG3 + CIA5/CCM1
ZNG3
CIA5/CCM1
Exclusively Zn induced
ZCP1
ZCP2
Zn and CO2 activated via Cia5
ZNG2
Cre09.g801069
Cia5-depedent low CO2 induced
Cre16.g692901
ZNG1
Neither Zn or CO2 activated
Cre03.g175700
Cre14.g613200
Repressed in low Zn
Cre16.g658000
Cre12.g536900
FPKM
dark
light
Zn
CO2
1
2
3
4
5

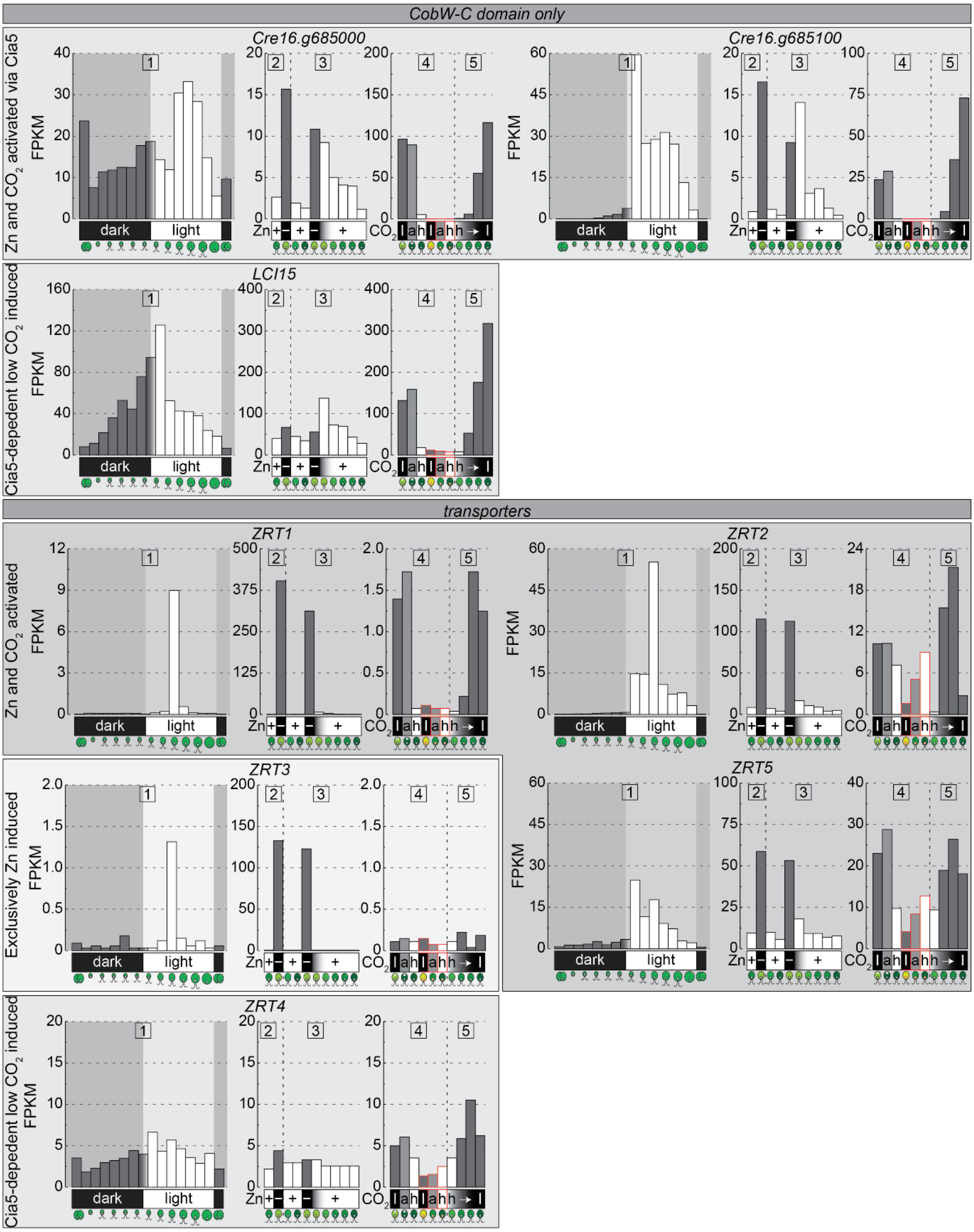


**Supplemental Figure 3. Transcript abundance of Zn importer and Zn chaperones in response to Zn deficiency, $CO_2$ supply and along the diurnal cycle.** Survey of transcript abundances of Zn transporters and candidate chaperones in published RNAseq datasets with varying Zn and $CO_2$ supply. Strenkert *et al.* (1) studied expression in phototrophically grown cultures along a diurnal cycle (12h dark/12h light), Malasarn *et al.* (2) analyzed transcripts in Zn- replete (+) and Zn-deficient (−) cultures, Hong Hermesdorf *et al.* (3) surveyed early exponential and early stationary Zn-replete cultures (+), as well as Zn-deficient (−) cultures and Zn resupply. Fang *et al.* (4) examined transcript abundances in cultures acclimated to high (+, 5%), air-level (=, 0.04%) and low $CO_2$ (−, 0.01%) in wildtype (black outline) and *cia5* mutants (red outline), while Brueggemann *et al.* (5) analyzed the transition from high (+, 5%) to very low $CO_2$ supply (−, 0.01%)

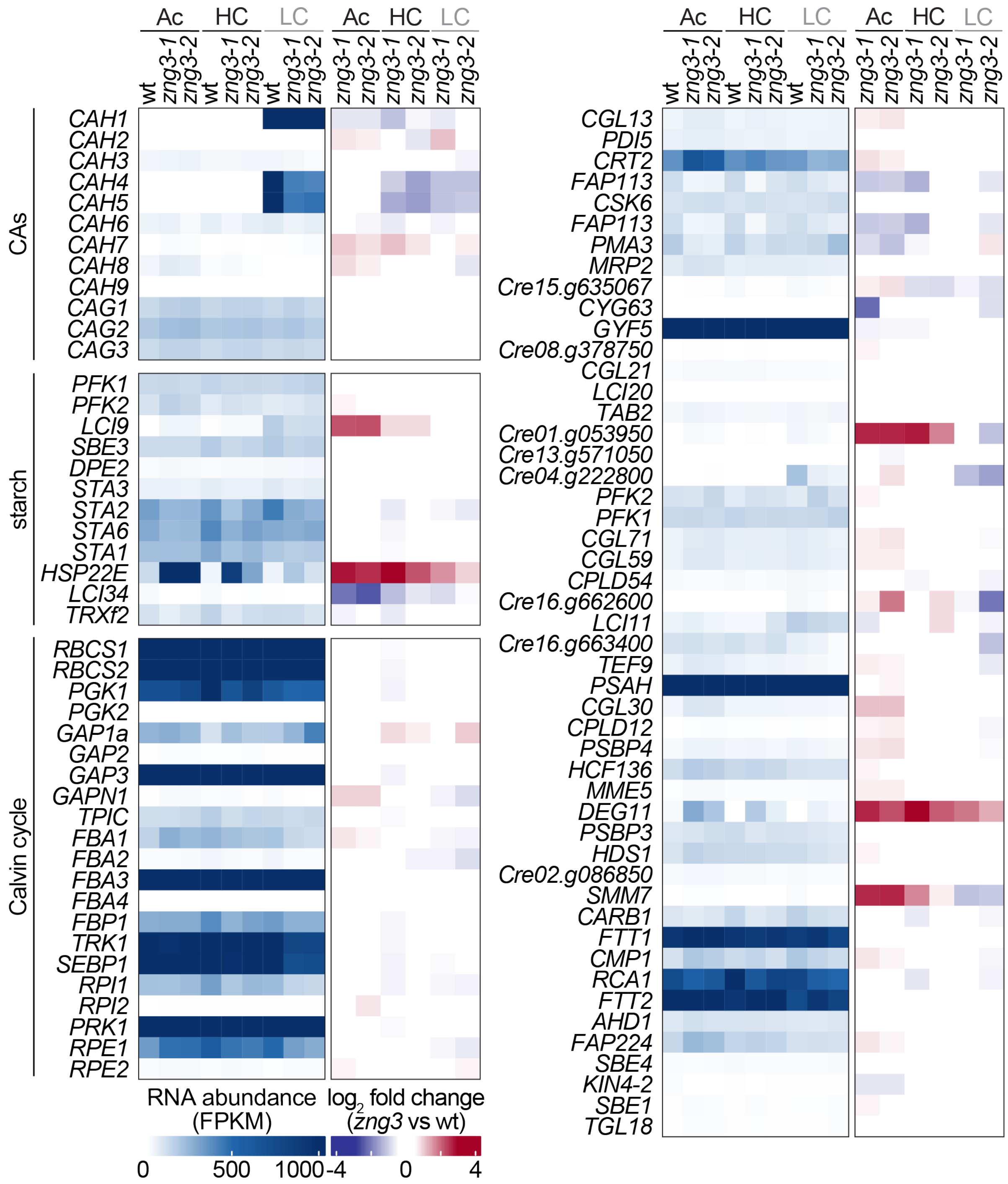


**Supplemental Figure 4. Expression of genes involved in the CCM and $CO_2$ assimilation.** Shown are transcript abundances and $log_2$ fold changes between *zng3* mutants and wildtype of genes encoding proteins involved in the CCM and $CO_2$ assimilation. The heatmap on the left highlights transcript abundances (fpkm), the heatmap on the right shows the $log_2$ fold changes between *zng3* and wt.

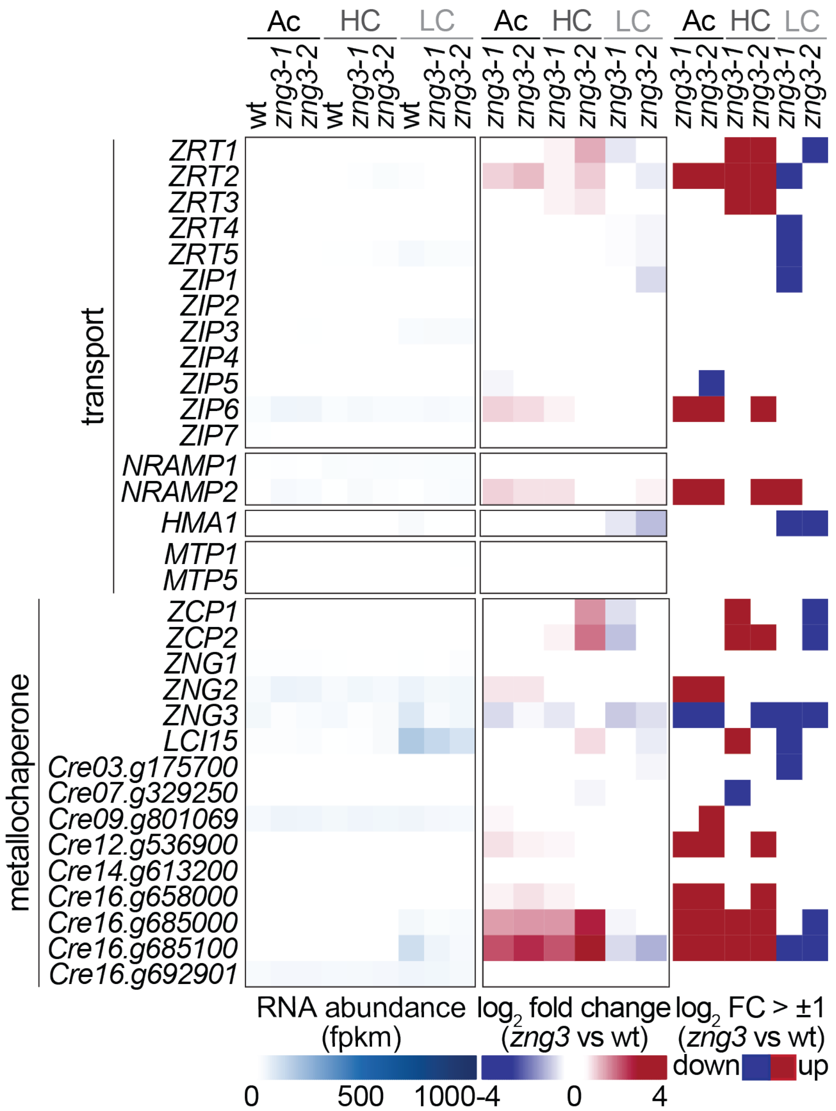


**Supplemental Figure 5. Expression of genes involved in Zn assimilation and distribution.** Shown are transcript abundances and $\log_2$ fold changes between *zng3* mutants and wildtype of genes encoding proteins involved in Zn assimilation and distribution. The heatmap on the left highlights transcript abundances (fpkm), the heatmap on the right shows the $\log_2$ fold changes between *zng3* and wt.

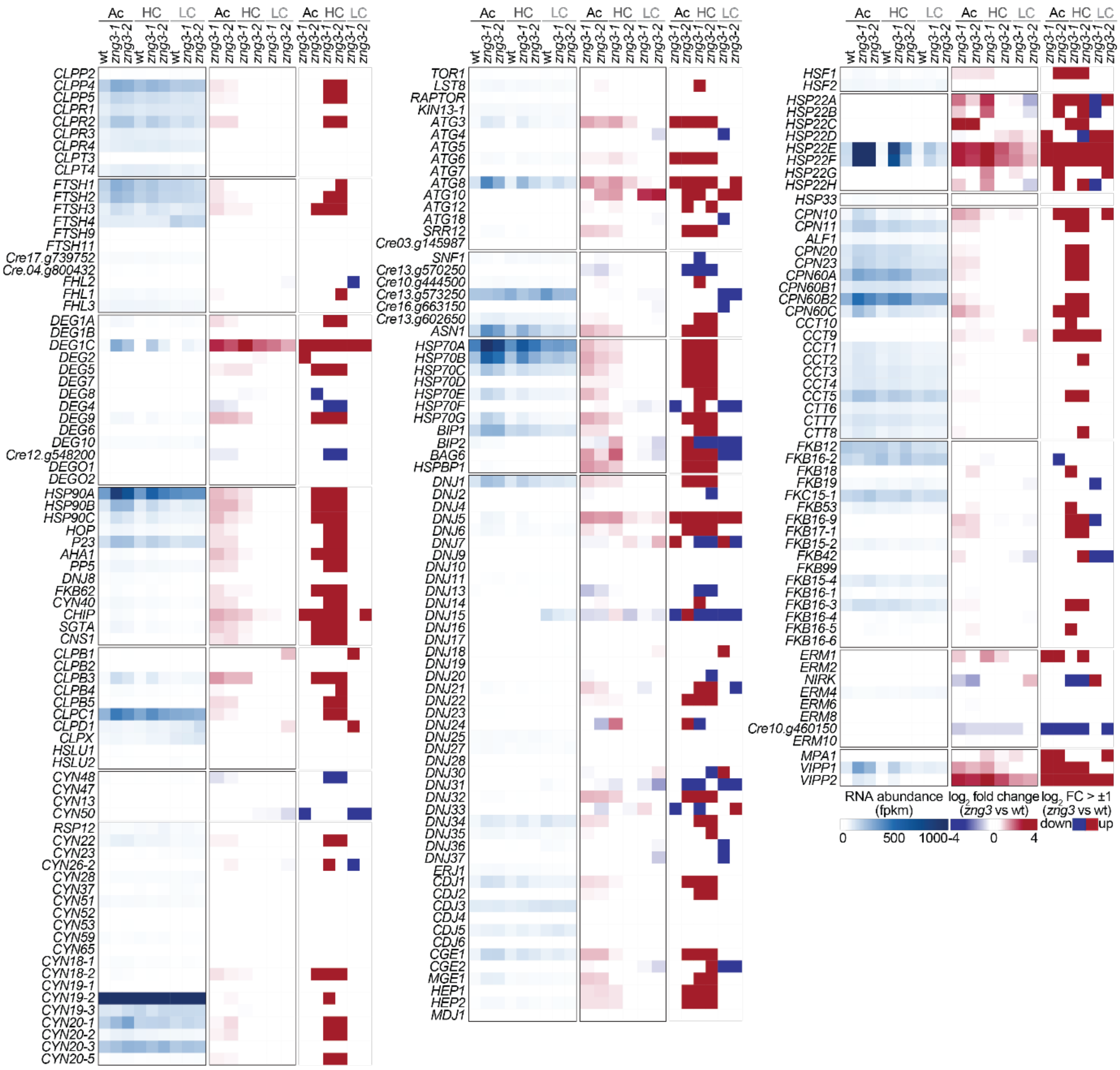


**Supplemental Figure 6. Expression of genes involved in protein quality control.** Shown are transcript abundances and $\log_2$ fold changes between *zng3* mutants and wildtype of genes encoding proteins involved in protein quality control. The heatmap on the left highlights transcript abundances (fpkm), the heatmap on the right shows the $\log_2$ fold changes between *zng3* and wt.
